# Neutral processes and high inter-annual turnover shape the assembly of soil bacterial communities in a Mediterranean watershed

**DOI:** 10.1101/542076

**Authors:** Myrto Tsiknia, Stilianos Fodelianakis, Nikolaos P. Nikolaidis, Nikolaos V. Paranychianakis

**Author notes:** Correspondence should be addressed to Nikolaos V. Paranychianakis, School of Environmental Engineering, Technical University of Crete, Polytechnioupolis 73100, Chania Greece.

## Abstract

There is a renewed interest in recent years on the ecological processes (stochastic vs selective) driving the assembly of microbial communities. Such information could potentially improve our understanding on ecosystem functioning and resilience to disturbances, ecosystem response to environmental shifts, and adoption of sustainable soil management practices. Herein, employing a suite of existing methodologies, we show that stochastic processes have an important role on the assembly of soil bacterial communities at a Mediterranean watershed. Moreover, we document that the relative contribution of assembly processes varies over the years. The observed intensification of stochastic processes was accompanied by a decrease in the contribution of variable selection in favor of homogeneous selection and dispersal and this trend was only marginally affected by land use (natural vs agricultural lands) or soil depth. Our study also revealed a high inter-annual turnover of soil microbial communities that was likely stimulated by the weak environmental selection and the prevailing environmental conditions (drying-wetting cycles) in Mediterranean landscapes, implying potential impacts on ecosystem functioning and our ability to predict soil response to environmental shifts. Using nitrogen mineralization rate (NMR) as a representative function we document highly variable NMR over the sampling years, land uses and soil depths and lack of significant associations with the monitored environmental variables and individual taxa. In summary, our study provides novel insights on the organization and functioning of microbial communities at Mediterranean ecosystems and sets directions towards a more advanced understanding of the relationships among environmental factors, microbial community structure, and ecosystem functioning that could contribute to sustainable management of these severely degraded ecosystems.

## Introduction

The structure of microbial communities has a critical role on the functioning of terrestrial ecosystems and the derived ecosystem services. Despite the accumulating knowledge in recent years on the mechanisms and factors that shape microbial communities, critical questions remain yet poorly addressed. A central issue regards the nature of the ecological processes (selective or stochastic) driving the assembly of microbial communities and whether shifts in the balance of assembly processes affect ecosystem functioning (Hanson et al 2012, Nemergut et al 2013, Zhou and Ning 2017). Such shifts might exert strong impacts on the response and resilience of ecosystems to disturbances imposed by anthropogenic activities (land use changes), physical factors (wildfires, wetting/drying cycles, droughts) or their combination (Guiot and Cramer 2016, Kefi et al 2007), but pertinent information remains yet scarce.

Several studies have revealed distinct geographical patterns of soil microbial communities that are typically attributed to variations in environmental factors (e.g., pH, organic matter), based on correlations or statistical modelling of the distribution of microbial taxa (Bru et al 2011, King et al 2010, Tsiknia et al 2014). Such relationships or models, however, do not necessarily represent cause-effect relationships since they account for only one of the possible aspects that could generate biogeographical patterns. It has been well accepted that geographical patterns of microorganisms are driven by both selective (e.g., environmental filtering, microbe-microbe interactions) and stochastic (dispersal, ecological drift, diversification) processes (Bahram et al 2016, Hanson et al 2012, Hao et al 2016, Lee et al 2017, Morrison-Whittle and Goddard 2015). Stochasticity is an inherent property of microbial communities assembled by neutral processes (Vellend et al 2014) according to the neutral theory (Hubbell (2001). The importance of neutral and selective processes in the assembly of microbial communities varies across and within ecosystems, depending on the succession stage, environmental conditions, or the disturbance type and severity (Dini-Andreote et al 2015, Ferrenberg et al 2013, Langenheder and Szekely 2011, Lee et al 2017, Ofiţeru et al 2010). Until recently, we lacked a methodological framework to quantify the contribution of assembly processes and their individual components (Stegen et al 2015) constraining efforts to interpret these findings with environmental parameters or process rates.

Compared to aquatic or engineered ecosystems (Gilbert et al 2012), most studies on terrestrial ecosystems report on snapshots of microbial community structure and, hence, their ability to provide reliable insights in longer timescales for the evolution of microbial communities and shifts in the importance of assembly processes remains questionable. Information on temporal dynamics of microbial communities can provide useful insights on variation sources, stability/resilience of microbial community to disturbances, and the interactions between taxa (Faust et al 2015) and could help to discern the impact of assembly processes on ecosystem functioning (Graham et al 2016, Stegen et al 2016).

To date, the processes shaping the assembly of soil microbial communities in Mediterranean landscapes remain poorly described. The distinct climatic conditions in Mediterranean ecosystems (e.g., drying/wetting cycles) and their high inter-annual variations might affect the relative contribution of assembly processes through their impacts on the individual mechanisms (selection, drift, dispersal, diversification). Earlier work at Koiliaris critical zone observatory (CZO) has revealed that microbial taxa, at the phylum level, are highly dispersed and environmental factors could account for a small proportion of the observed variance (Tsiknia et al 2014). This finding is indicative of a potentially important role of neutral processes in the assembly of soil microbial communities in this Mediterranean ecosystem. In addition, in absence of major disturbance events (land use changes, drought) in recent years, we hypothesize that soil microbial community have reached a stable state (Fukami 2015) with slight variation over the years.

To shed light on the processes regulating the assembly of microbial communities in Mediterranean ecosystems, we performed a triennial sampling that included sites with natural vegetation and agricultural fields sampled at two soil depths. We employed a suite of available methodologies, including the Sloan’s neutral community model (SNCM) (Sloan et al 2007), βNTI metric (Stegen et al 2013), and variation partitioning analysis (VPA) (Peres-Neto et al 2006) to disentangle the importance of neutral and selective processes on microbial community assembly and its inter-annual variation. Subsequently, we quantified the relative contribution of assembly processes and the underlying mechanisms using the βNTI and the modified Raup-Crick (RC_Bray_) metrics (Stegen et al 2015). This approach allowed us to address inherent limitations of the existing methods and to reliably infer for the underlying mechanisms of assembly and their importance (Zhou and Ning 2017). Finally, we interpret variations in the relative contribution of assembly processes over the years and land uses with a central process of ecosystem functioning, the net nitrogen mineralization rate (NMR), to identify potential links between the importance of assembly processes.

## Materials and Methods

The Koiliaris CZO watershed is located 25 km on the western part of Crete, Greece (005-12-489E, 039-22-112N). The watershed occupies an area of approximately 130 km^2^ with the altitude to range from 0 to 2100 m. In this study three soil samplings were performed on May 10-12, 2012, May 10-13, 2013, and May 12-14, 2014. Representative fields to capture the basic features of the watershed (land, use, crops, latitude) were selected (Supplementary Figure S1). Information on the variation of climatic conditions in the watershed during the study period is summarized in Supplementary Figure S2. Twenty-two composite (three replicates/site) samples were collected from two soil depths; 0-15 (all the years) and 15-30 cm (in 2013 and 2014 samplings). Samples were passed through a 2mm sieve and were stored on ice packs at 4 °C until their transport to the laboratory. Soil physical (pH, texture), chemical (TOC, TN, NH_4_^+^-N, NO_3_^-^-N) and biochemical (NMR) analyses were performed using the methods described in (Tsiknia et al 2014) and are summarized in Supplementary Table S1. DNA was extracted with the PowerSoil^®^ DNA isolation kit according to the manufacturer’sinstructions.

### Amplification and sequencing of the 16S rRNA gene and data processing

The universal prokaryotic primer set 515f/806r was used to amplify the V4 hypervariable region of the 16S rRNA gene. Amplicon sequencing was performed at a MiSeq Illumina platform at Research & Testing labs (Austin, TX, USA). The raw sequences were analyzed using a combination of the UPARSE (Edgar 2013) and the QIIME (Caporaso et al 2010) pipelines. Briefly, raw forward and reverse reads for each sample were assembled into paired-end reads considering a minimum overlapping of 50 nucleotides and a maximum of one mismatch. The paired reads were then quality filtered, and the primer sequences were removed. The UPARSE pipeline was used for OTUs construction and chimeras were removed using both de-novo (Edgar *et al.* 2011) and reference-based (Reddy et al 2015) methodologies. Taxonomy was assigned to the representative sequences of OTUs in QIIME v. 1.8 and singletons were removed from the resulted OTU table before further analysis. Finally, the phylogenetic tree was constructed with the FastTree algorithm using default parameters (Price et al 2010). Raw sequences have been deposited in the ENA European Read Archive with accession number PRJEB22862.

### α- and β-diversity and statistical analyses

Alpha diversity was determined by various indices that estimate community’s richness (observed OTUs and Chao1), total diversity (Shannon, InvSimpson, Fisher’s α), evenness (Pielou’s J), and phylogenetic diversity (Faith’s PD), either with the QIIME pipeline or with the R software using the *vegan* (Oksanen et al 2013) and *phyloseq* (McMurdie and Holmes 2013) packages. Non-parametric factorial ANOVA with mixed-effect models was performed with the ARTool package (Wobbrock et al 2011) and was used to investigate the main effects and the interactions of the sampling year, soil depth and land use (aggregated in agricultural lands vs natural ecosystems) on α-diversity indices and environmental variables monitored in the present study. Year, soil depth, and land use were set as fixed-effect factors and sampling site as random effect.

Spearman correlation was used to identify significant links between environmental parameters and α-diversity indices, annually. To quantify the annual variation of soil parameters, the coefficient of variation (temporal CV %) was calculated from the three replicate samples of each site. Permutational multivariate analysis of variance (PERMANOVA; *adonis()* function, with 999 permutations) was applied separately for each year to both the Bray-Curtis dissimilarity matrix and weighted UniFrac distance matrix, to assess the effect of depth and land use on β-diversity. Distance based redundancy analysis (db-RDA) was applied to visualize β-diversity patterns, with both Bray-Curtis dissimilarity matrix and the weighted UniFrac distance matrix.

### Determining community assembly dynamics

The importance of microbial community assembly processes was evaluated by the Sloan’s neutral community model (SNCM) for prokaryotes (Sloan et al 2007) and null modeling (Stegen et al 2012). The SNCM has been evolved from the Hubbell’s neutral community theory (Hubbell 2001). The relationship of the mean relative abundance and frequency of occurrence of a taxon belonging to a neutral community is described by a beta distribution. Significant deviations from the SNCM, e.g., more- or less-abundant taxa than predicted, imply communities that are assembled by selective processes or deviations from the mean coefficient of immigration (m). A binomial distribution was also fitted to the above relationship to test whether microbial communities were derived by random sampling. The significance of fit of the neutral and binomial models was compared with the Akaike information criterion (AIC). To fit the SNCM to the data and to estimate model’s goodness-of-fit and AIC, we adopted the methodology and the R code developed by Burns et al (2016). Afterwards, the OTUs were separated into three distinct groups according to the model prediction; OTUs which were as abundant as the model predicted (neutral OTUs), and OTUs which occurred with higher or lower abundance than it was predicted (non-neutral OTUs).

Additionally, we quantified the phylogenetic dispersion of the whole microbial communities and the groups separated by the SNCM to obtain deeper insights on the underlying mechanisms of community assembly. The β-nearest taxon index (βNTI) was used to assess the deviation between the observed and the null β-mean nearest taxon distance (βMNTD) (after 999 randomizations). The latter represents the phylogenetic dispersion of a community dominated by stochastic processes, while any deviation underlines the importance and the homogeneity of selective processes. The null distribution of βMNTD was calculated from random subsampling (999 times) and subsequent pairwise comparisons of the phylogenetic tree of the whole metacommunity, while the observed βMNTD was calculated from pairwise sample comparisons. To confirm the suitability of βMNTD as a β-phylogenetic diversity metric, we first performed a phylogenetic signal test (Stegen et al 2013) to examine whether the ‘habitat preferences’ of OTUs are more similar at short or large phylogenetic distances. We first calculated the ‘habitat preferences’ of the OTUs (based on pH, soil moisture, NO_3_-N, TOC, C/N ratio and clay content) by finding the niche optima of each OTU for each parameter (Fodelianakis et al 2017) and creating a similarity matrix from the resulting table (function vegdist of the vegan package in R, method=“euclidean”). We then created a phylogenetic distance matrix from the phylogenetic tree of the OTUs (function cophenetic of the stats R package) and examined the correlation with the environmental matrix at increasing phylogenetic distance classes (function mantel.correlog of the vegan package in R, Pearson correlation, 999 permutations). We found significant correlations at short phylogenetic distances (Supplementary Figure S3), indicating the use of β-MNTD as a phylogenetic metric. To estimate the βNTI, the null βMNTD was calculated by random subsampling of the phylogenetic tree, and the observed βMNTD was calculated from all pairwise comparisons. For these analyses, we used the “ses.mntd()” function of the picante package (Kembel et al. 2010).

To quantify the relative contribution of the underlying mechanisms (e.g. selection, drift, dispersal) we adopted the framework developed by (Stegen et al 2015). Shortly, the variation in phylogenetic and taxonomic diversity is estimated with the βNTI, and RCBray metrics respectively. Homogeneous selection results in phylogenetically similar communities and, hence, the percentage of homogeneous selection can be estimated as the fraction of pairwise comparisons with a βNTI value < −2. In accordance, heterogeneous selection is quantified as the fraction of pairwise comparisons with βNTI values > 2. Then, the RCBray metric is employed to further partition the pairwise comparisons with an βNTI values < |2|. Since homogenizing dispersal results in taxonomically similar microbial communities, its relative importance is quantified as the fraction of the pairwise comparisons with βNTI and RC_Bray_ values < |2| and < −0.95 respectively. Dispersal limitation leads to communities less similar in taxonomy and can be quantified as the fraction of the pairwise comparisons with βNTI and RC_Bray_ values < |2|and > 0.95 respectively. Finally, values of βNTI and RC_Bray_ metrics < |2| and > |0.95| respectively correspond to the “undominated” fraction by the weak influence of selection, dispersal, diversification, and drift.

The contribution of assembly processes can be alternatively estimated by multivariate analysis. Despite its inherent limitations (Hanson et al 2012), certain types of multivariate analysis, like the variation partitioning analysis (VPA), can provide improved estimations of the importance of selection. VPA based on redundancy analysis (RDA) of distances (Peres-Neto et al 2006) was performed to investigate the influence of soil variables, spatial structure, and their interaction on β-diversity patterns over the years, in different land uses and soil depths. To perform the analysis, all the spatial scales that could be perceived in the study area were determined by the Principal Coordinates of Neighbor Matrices (PCNM) method (Borcard and Legendre 2002) as it was described in Tsiknia et al (2015). The positive PCNM vectors constituted the spatial matrix that was used in VPA. In the following step, the most significant variables were identified with forward model selection, using the *ordiR2step()* function of the vegan package, separately for the soil variables and for the spatial dataset, and were subsequently used for VPA using *varpart()* function. All the employed methodologies (SNCM, null modeling, VPA) were run separately for each year, land use, and soil depth to investigate the influence of these factors on the assembly processes of soil microbial community.

## Results

### Microbial community structure and diversity

Detailed information on the taxonomy of soil microbial communities is presented in Supplementary Figure S4 and text. Mixed-effects ANOVA identified the sampling year as the only factor that affected significantly α-diversity (Supplementary Table S2). Richness (observed OTUs and Chao1) and diversity (InvSimpson) estimators decreased in 2014 compared to 2012 and 2013 (Supplementary Table S3). α-diversity metrics were correlated with several soil variables, but these relationships were inconsistent over the years (Supplementary Table S4). db-RDA of weighted UniFrac distance identified pH, clay content, soil moisture, TOC content, and C:N ratio as the most important drivers of β-diversity, explaining up to 18.6% of total variance (Supplementary Figure S5A). Similar results were obtained with the Bray-Curtis dissimilarity (Supplementary Figure S5B). PERMANOVA on both distances revealed a strong influence of the main factors on β-diversity patterns, with the sampling year to explain a greater proportion of the observed variance than land use and soil depth (Supplementary Table S5).

### Importance of assembly processes over the years, soil depths and land uses

The high coefficient of determination of the SNCM (Figure 1) suggests that neutral processes have an important role in microbial community assembly at Koiliaris CZO. Application of the SNCM separately for each sampling year provided support for shifts in the importance of neutral processes over year (Figure 1a-c) as it could be inferred by the increases of the coefficients of determination and immigration rate (m). The fit of SNCM did not diverge substantially with soil depth or land use (Supplementary Figure S6 and S7). In all years, soil depths, and land uses the SNCM outperformed binomial distribution (Supplementary Table S6). Regarding the OTUs partitioned above or below the confidence intervals of the model’s prediction, they likely correspond to taxa that are either selected for/against by the prevailing environmental conditions or are dispersal limited (Burns et al 2016). NMDS analysis of Bray-Curtis dissimilarities and Unifrac distances revealed that the neutral, less frequent than predicted and more frequent than predicted OTUs formed clearly separated clusters (Supplementary Figure S8) pointing out to divergent mechanisms of assembly.

**Figure 1.**
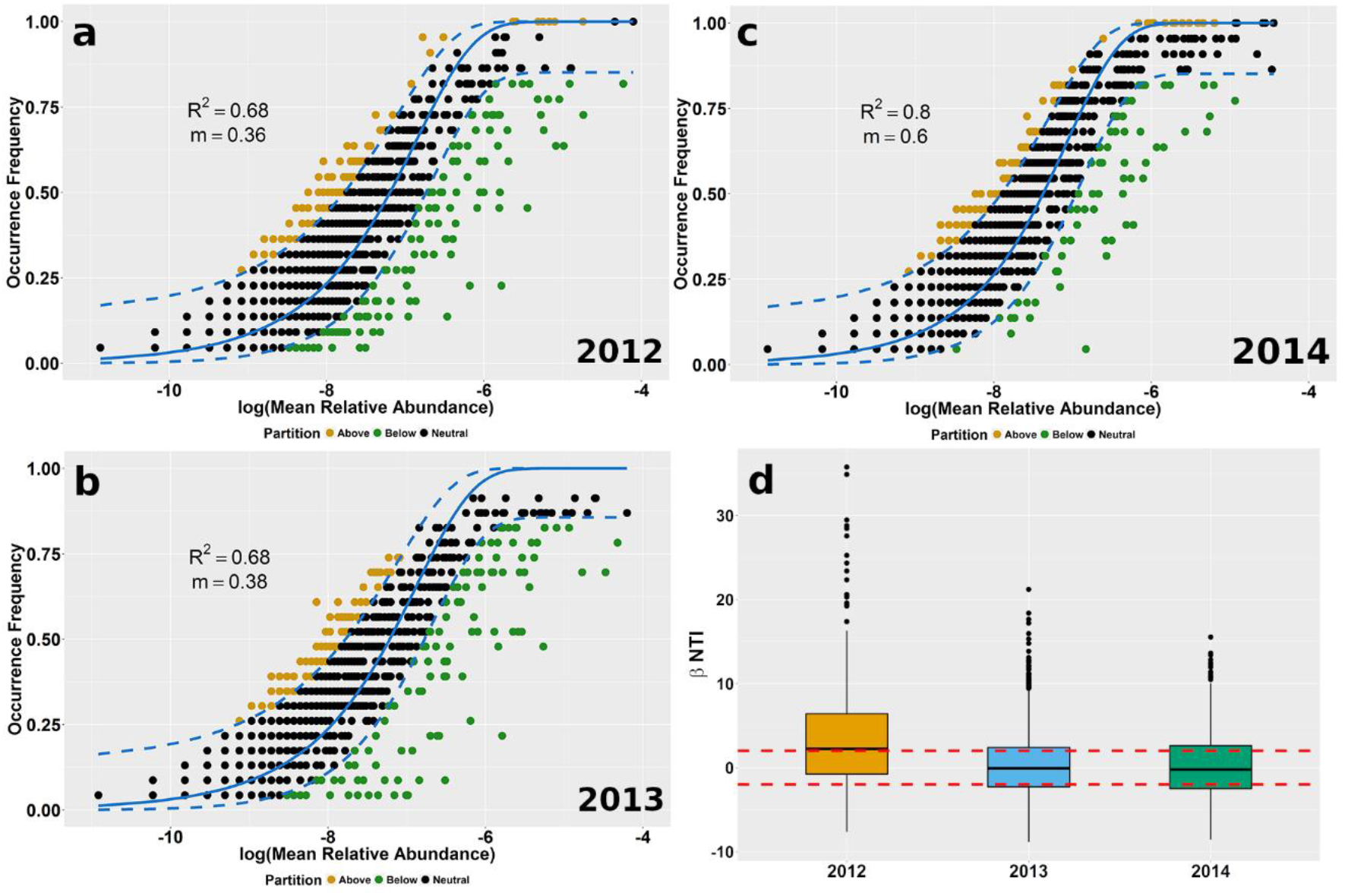
Importance of microbial community assembly processes over the sampling period (2012-2014). Fit of Sloan Neutral Community Model for Prokaryotes in soil bacterial communities. In each plot, the model fit (R2) and immigration rate (m) are shown for (a) 2012, (b) 2013, and (c) 2014 samplings. OTUs that occurred more frequently than expected are shown in yellow, while OTUs that occurred less frequently than expected are shown in green. Dashed lines represent the 95% confidence intervals of the model prediction (solid line). (d) Box plots of βNTI distributions over the sampling period showing the median (thick black line), the first quartile (lower box bound), the third quartile (upper box bound), and outliers (black circles). Horizontal red dashed lines indicate the upper and lower significance thresholds of βNTI at +2 and −2, respectively.

To further confirm the importance of neutral processes, we used an alternative metric, the βNTI. The mean βNTI decreased from 3.52 in 2012, a value indicative of the prevalence of selective processes, to −0.15 in 2013, and to −0.17 in 2014 providing further support that neutral processes had an important role in microbial community assembly (Figure 1d), confirming partially the findings of the SNCM. The βNTI, however, revealed a more important role of neutral processes in microbial community assembly in natural ecosystems than agricultural lands, whereas, similar conclusions were reached for soil depth (Supplementary Figure S9).

The composition of the whole community and the communities separated by the SNCM changed substantially over the years. We identified 2646 out of the 8227 OTUs (32.2%) that were present all the sampling years (Figure 2), while significant turnover was also observed for the neutral community, and the less frequent, and more frequent than expected OTUs (Supplementary Figure S10).

**Figure 2.**
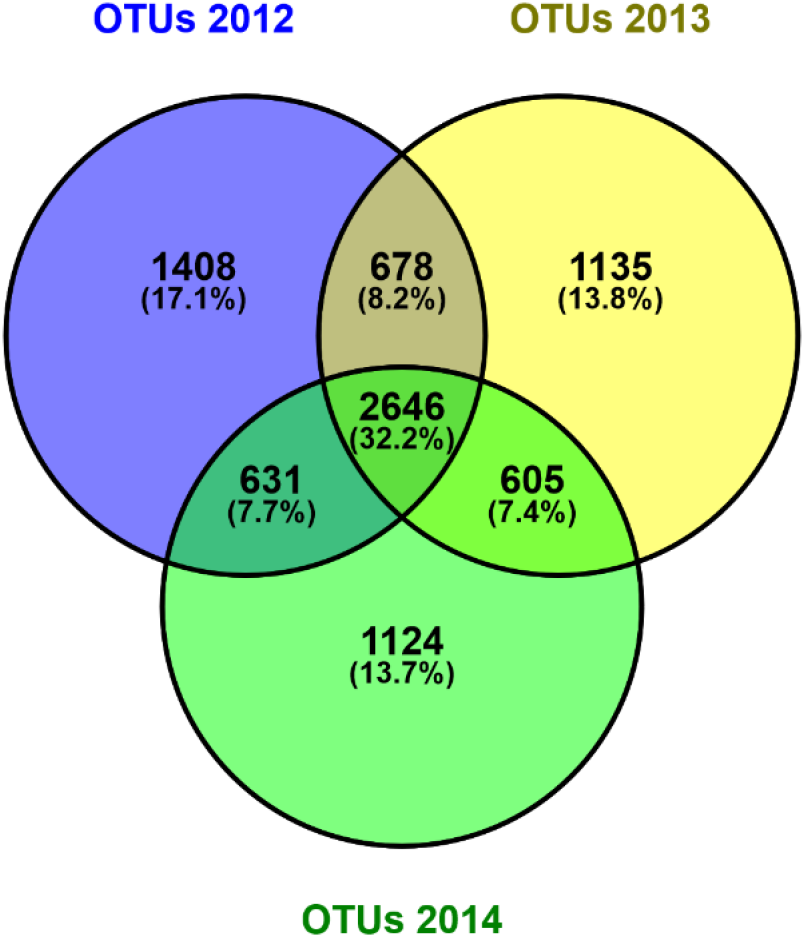
Interannual turnover of soil microbial communities. Venn diagrams of the common OTUs for the three sampling years

### Relative contribution of assembly processes

In the following step, we employed VPA and RCBray methodologies to obtain estimations of the relative contribution of neutral and selective processes on β-diversity patterns of soil bacterial community. The proportion of β-diversity that was purely explained by soil variables decreased from 14% in 2012 to 4% in 2013 and to 2% in 2014 (Figure 3; Supplementary Table S7), while the corresponding proportion of the spatial variables increased from 5% in 2012 to 14% in 2013 and then decreased to 8% in 2014 (Figure 3). The shared variation of the soil and spatial variables remained low throughout the study period. The proportion of unexplained variance followed an increasing trend during the study period (Figure 3). This trend remained unchanged independently of soil depth and land use, with most of the variance (71-80%) remaining unexplained (Supplementary Figure S11), assigning a dominant role on stochastic processes in community assembly. In accordance, the distance-decay plots of Bray-Curtis similarity followed a weak decay pattern all the years (Supplementary Figure S12).

**Figure 3.**
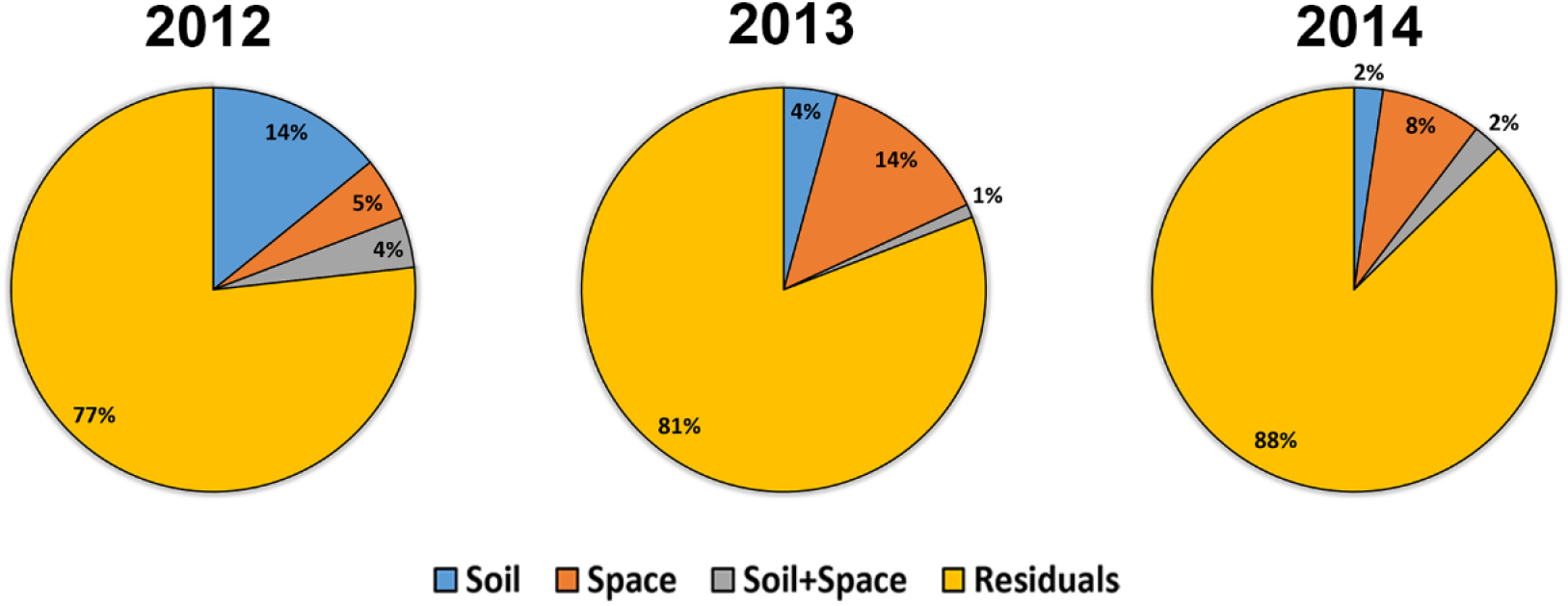
Variation partitioning analysis over the years of the effects of soil and spatial variables on β-diversity. The partitions indicate the percentage of variation explained by the soil variables (Soil), the space (Space), the shared effect of soil and space (Soil+Space), and the unexplained variation (residuals).

The RC_Bray_ framework provided quite different estimations of the relative contribution of neutral and selective processes and their individual mechanisms. More specifically, the relative contribution of selective processes (homogeneous and variable) declined from 68% in 2012 to 50% in 2014 (Table 1). Interestingly, this decrease was accompanied by a shift of the contribution of variable selection in favor of homogeneous selection. The observed pattern did not change substantially with soil depth (Supplementary Table S8), but significant differentiations were found between land uses with the relative contribution of variable selection to be greater in agricultural lands compared to natural ecosystems (Supplementary Table S8).

**Table 1.**
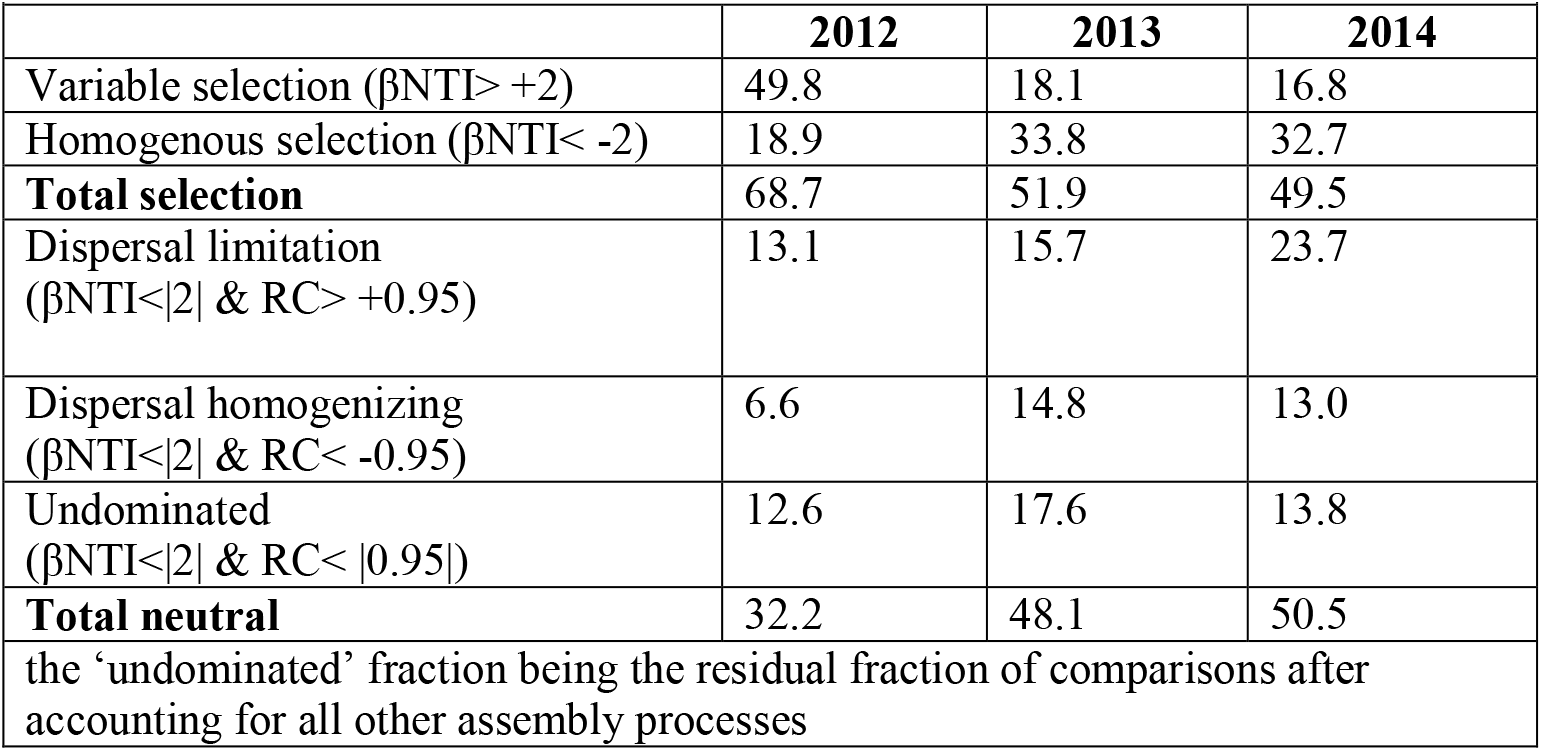
Relative contribution (%) of individual mechanisms of assembly (dispersal, selection, drift) on soil microbial communities during the 2012-2014 samplings. The estimation of the contribution is based on the framework of Stegen et al (2015).

### Links between Assembly Processes and Functional Process rates

To test for links between functional process rates and relative contribution of assembly processes, we monitored NMR. The NMR showed high variation between sites and years, particularly in the upper soil layer (Supplementary Figure S13) and was not correlated significantly with any of the soil variables monitored in the study. To obtain insights on the potential links between shifts in microbial community structure and NMR, we employed the βNTI metric. Shifts in βNTI are driven principally by variations in the selective environment and to a lesser extent by neutral processes, like dispersal (Stegen et al 2016). Despite the variation of NMR between sites, no significant relationship was found with βNTI and that effect was independent of year, soil depth, or land use (Figure 4; Supplementary Figures 14 and 15). Regarding the relationships of NMR with individual taxa, a few significant correlations were set and only at a high taxonomic level (Phylum or Class) (Supplementary Table S9). These relationships were, however, highly variable between land uses, sampling years and soil depths implying probably spurious and/or indirect relationships. These findings strengthen the assumption that stochastic processes have a more important role than environmental selection in shaping soil microbial communities in that Mediterranean watershed. That, in turn, uncoupled processes rates from environmental variables and shifts in soil microbial community structure.

**Figure 4.**
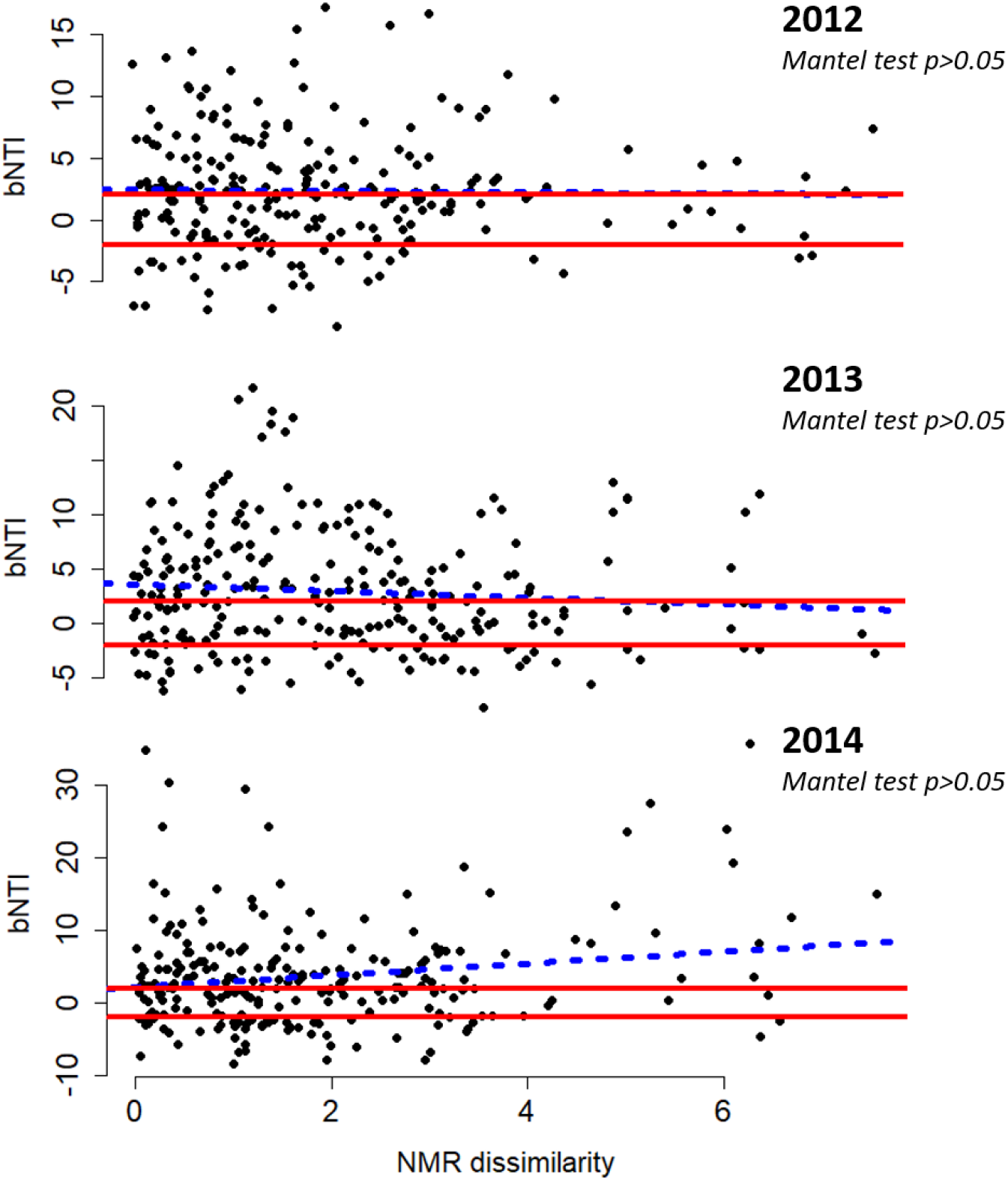
The relationship between βNTI and NMR dissimilarity (Euclidean distance) for each sampling year. The regression slopes of the linear relationships, based on Gaussian generalized model, are shown with blue dashed lines (non-significant; Mantel test; 9999 permutations; p> 0.05). The solid red line indicates the upper and lower significance thresholds at βNTI=+2 and −2, respectively.

## Discussion

Employing different conceptual approaches (neutral theory, null modeling, VPA), we provide convergent support that neutral processes have an important role in the assembly of soil microbial communities in a typical Mediterranean watershed. This finding is consistent with earlier work in Koiliaris CZO that has revealed weak links between environmental variables and the abundance of bacterial phyla (Tsiknia et al 2014).

The high coefficients of determination and immigration estimated by the SNCM provide evidence for an important role of neutral processes (Burns et al., 2016), that is well aligned to the findings obtained by VPA. Both methodologies, however, suffer from biased estimations in favor of neutral processes. In the case of SNCM this arises from the indirect estimation of parameters by optimizing model fitting to microbial community data (Sloan et al 2007). For instance, empirical studies in specialized environments (e.g., bioreactors) have assigned an unexpectedly high importance to neutral processes (Ayarza and Erijman 2011, Ofiţeru et al 2010, Zhou et al 2013). In the case of VPA the overestimation of neutral processes arises from the non-consideration of unmeasured environmental variables or biotic interactions (Smith & Lundholm, 2010). By contrast, the framework developed by (Stegen et al 2015) revealed a milder contribution of stochastic processes that eventually reached and exceeded that of selective processes in the 2^nd^ and 3^rd^ sampling years, respectively. This framework has been used in several studies providing realistic findings.

### Temporal shifts in the relative contribution of assembly processes

To date, temporal shifts in the importance of microbial assembly processes have been documented intra-seasonally in soils recovering from fire disturbance (Ferrenberg et al 2013) or in soils at different stages of succession (Dini-Andreote et al 2015). This is the first study to document strong inter-annual variations in the relative contribution of assembly processes in the soils of a Mediterranean watershed. Since the watershed has not recently experienced severe disturbances (e.g., fires, land use change, droughts), the observed shifts have likely arisen by variations of climatic factors which, in turn, have affected the strength of individual mechanisms of assembly (e.g., dispersal, drift, selection) in the soils. A similar temporal trend has been reported in lakes, where the importance of neutral processes increased over a triennial period and that trend was accompanied by an increase in immigration coefficient (Roguet et al 2015).

In addition, the βNTI and RCBray framework (Stegen et al 2015) allowed us to get insights into the shifts of individual mechanisms of assembly. The remarkable decrease in the contribution of variable selection in the 2^nd^ and 3^rd^ year, partially in favor of homogenizing selection, likely imply intensification of the influence of environmental variables (e.g., temperature and/or soil moisture) that favor the latter. Such an effect could have outweighed the influence of spatial variations in soil properties (texture, pH, SOM) or management practices that are responsible for variable selection (Rousk et al 2010). This explanation is consistent with the greater contribution of homogeneous selection in natural ecosystems that are concentrated at the upper part of the watershed (Figure S1) and are characterized by lower heterogeneity of environmental and biotic factors (e.g., vegetation type, soil properties) compared to agricultural fields. The role of dispersal in community assembly remains, however, obscure due to the quite variable estimations obtained by the different methodologies. A strong influence of dispersal, as it could be inferred by the SNCM, would have overcome the effect of weak selection and have resulted in homogenized microbial communities (Evans et al 2017, Leibold et al 2004). The milder dispersal rates that have been estimated by the Stegen et al. (2015) framework and acted bidirectionally (dispersal limitation and homogenizing dispersal; Table 1) appear to provide more realistic estimations considering the observed spatial patterns of microbial communities as well as the geomorphology of the watershed (Nerantzaki et al 2015). The high inter-annual turnover of microbial communities provides some evidence that the sources of dispersal extend beyond the boundaries of watershed. Wind erosion and African dust events that commonly occur in Mediterranean basin (Polymenakou et al 2008) could be important external sources of bacteria with a strong impact on the structure of soil bacterial communities (Ma et al 2017).

### Interannual turnover of soil microbial communities

Microbial communities tend to approach a stable state after ecosystem succession and in absence of severe disturbance events (Chase 2003, Fukami 2015), but such a trend was not observed at Koiliaris CZO. The seasonal shifts in precipitation, temperature and availability of resources (birch effect) (Placella et al 2012) during the summertime likely act as disturbing agents of soil microbial communities by decreasing the abundance, diversity, and activity of microbial communities (Ma et al 2015, Maestre et al 2015) in Mediterranean landscapes. The re-wetting of soils in autumn relieves the imposed constraints to microorganism growth and activity initiating, thus, new rounds of microbial community assembly. Considering the stochastic nature of immigration (magnitude and timing) and ecological drift, they likely had a prominent effect on the divergence of soil microbial community from year to year as has been reported in earlier works (Fukami 2015). Particularly in absence of strong selection filters, as it has been found in Koiliaris CZO, the influence of immigration and ecological drift is strengthened (Nemergut et al 2013) and might have stimulated the observed high inter-annual compositional turnover of microbial communities.

### Assembly processes and ecosystem functioning

A fundamental question that arises from the inter-annual turnover of microbial communities and the importance of neutral processes of assembly regards their impacts on ecosystem functioning. Dominance of neutral processes in the hyporheic zone of a riverine system resulted in highly variable rates of respiration (Stegen et al 2016). Likewise, divergence of microbial communities in pilot bioreactors, driven by neutral processes, differentiated strongly the performance of bioreactors (Zhou et al 2013). These findings are consistent with the highly variable rates of NMR that were measured in this study over the years and the lack of significant correlation between NMR and any of the soil variables monitored, independently of land use or soil depth. In accordance, environmental variables could explain only a small proportion of the observed variance in litter decomposition rate which however, was significantly linked to soil microbial biomass (Bradford et al 2017). A few significant relationships between NMR and microbial taxa were set in this study mainly at high taxonomic level. These relationships, however, were quite variable over the years and between land uses and soil depths likely reflecting the high turnover of microbial communities and their spatial variability. These findings challenge earlier suggestions that inclusion of information for microbial communities assembled by neutral processes is necessary to obtain improved predictions of ecosystem functioning (Ferrenberg et al 2013).

By providing quantifications of the relative contribution of neutral processes in microbial community assembly, we advance existing knowledge and show that even mild contribution of neutral processes, up to 30%, can generate high variability in process rates increasing the uncertainty of model simulations. These findings have profound implications on the adoption of appropriate management practices and on their effectiveness in the severely degraded soils of the Mediterranean watersheds (Moraetis et al 2015). Despite the growing concern in maintaining/improving soil health by managing microbial communities (Wallenstein 2017, Wubs et al 2016), the great compositional turnover of microbial communities questions the success of this practice in Mediterranean ecosystems. A more comprehensive understanding is urgently needed on the feedbacks between environmental drivers and assembly processes, the critical thresholds of neutral processes that increase variation of process rates, and the impact of agricultural practices on the balance of assembly processes to improve the potential for “engineering” soil microbial communities and to maintain them in the long-term.

## Conclusions

The present study advances our understanding on the processes driving the assembly of microbial communities in Mediterranean ecosystems. The data provides support that neutral processes have an important role on the assembly of bacterial communities with variable, however, contribution over the years. Dispersal, either limiting or homogenizing, has been identified as a strong driver of microbial community assembly. The importance of neutral processes in combination with environmental conditions (weak environmental selection) are likely the main drivers that stimulate the high inter-annual turnover of soil microbial communities. The highly variable rates of NMR between years, sites, and land uses, used as a proxy of soil functioning, were not correlated with any of the environmental variables monitored in the study and likely reflect the stochasticity in microbial community assembly. Considering the accumulating evidence for intense climatic changes in the Mediterranean (drought frequency, temperature increase, precipitation patterns), these findings raise critical questions about soil functioning, like their response to environmental shifts, their resilience to disturbances and our ability to reliably simulate soil functioning with conventional biogeochemical models that only implicitly consider the role of microbial communities. However, inclusion of microbial communities in models appears to be a challenging task since strong relationships were only set at high taxonomic levels and theses relationships changed between years, land use and soil depths. These findings underline the need for additional studies to advance our understanding on the complex interactions between microbial community assembly processes, environmental variables and microbial community properties and on their effect on key ecological processes (e.g., C and N cycling) providing a basis for the development of soil adaptation practices to secure the sustainable use of soils and maintain their productivity in the long-term.

## Author contribution

NVP and NN concepted and designed the study. NN contributed consumables. MT, NVP and SF performed the samplings. MT performed soil chemical and biochemical analyses. MT performed statistical analysis. MT and SF performed the bioinformatic analyses. NVP wrote the manuscript with contributions from MT and SF. All authors contributed to and approved the revisions.

## Conflict of Interest

The authors declare no conflict of interest.

## Acknowledgements

Funding was provided from the European Commission FP7 Collaborative Project Soil Transformations in European Catchments (SoilTrEC) (Grant Agreement No. 244118) and resources from the Technical University of Crete.

## References

Ayarza JM, Erijman L (2011). Balance of neutral and deterministic components in the dynamics of activated sludge floc assembly. Microb Ecol 61: 486–495.

Bahram M, Kohout P, Anslan S, Harend H, Abarenkov K, Tedersoo L (2016). Stochastic distribution of small soil eukaryotes resulting from high dispersal and drift in a local environment. ISME J 10: 885–896.

Borcard D, Legendre P (2002). All-scale spatial analysis of ecological data by means of principal coordinates of neighbour matrices. Ecol Model 153: 51–68.

Bradford MA, Veen GF, Bonis A, Bradford EM, Classen AT, Cornelissen JHC et al (2017). A test of the hierarchical model of litter decomposition. Nature Ecology & Evolution 1: 1836–1845.

Bru D, Ramette A, Saby NP, Dequiedt S, Ranjard L, Jolivet C et al (2011). Determinants of the distribution of nitrogen-cycling microbial communities at the landscape scale. ISME J 5: 532–542.

Burns AR, Stephens WZ, Stagaman K, Wong S, Rawls JF, Guillemin K et al (2016). Contribution of neutral processes to the assembly of gut microbial communities in the zebrafish over host development. ISME J 10: 655–664.

Caporaso JG, Kuczynski J, Stombaugh J, Bittinger K, Bushman FD, Costello EK et al (2010). QIIME allows analysis of high-throughput community sequencing data. Nat Methods 7: 335–336.

Chase JM (2003). Community assembly: when should history matter? Oecologia 136: 489–498.

Dini-Andreote F, Stegen JC, van Elsas JD, Salles JF (2015). Disentangling mechanisms that mediate the balance between stochastic and deterministic processes in microbial succession. Proc Natl Acad Sci USA 112: E1326–E1332.

Edgar RC (2013). UPARSE: highly accurate OTU sequences from microbial amplicon reads. Nat Meth 10: 996–998.

Evans S, Martiny JBH, Allison SD (2017). Effects of dispersal and selection on stochastic assembly in microbial communities. ISME J 11: 176–185.

Faust K, Lahti L, Gonze D, de Vos WM, Raes J (2015). Metagenomics meets time series analysis: unraveling microbial community dynamics. Current Opinion in Microbiology 25: 56–66.

Ferrenberg S, O’Neill SP, Knelman JE, Todd B, Duggan S, Bradley D et al (2013). Changes in assembly processes in soil bacterial communities following a wildfire disturbance. ISME J 7: 1102–1111.

Fodelianakis S, Moustakas A, Papageorgiou N, Manoli O, Tsikopoulou I, Michoud G et al (2017). Modified niche optima and breadths explain the historical contingency of bacterial community responses to eutrophication in coastal sediments. Molecular Ecology 26: 2006–2018.

Fukami T (2015). Historical Contingency in Community Assembly: Integrating Niches, Species Pools, and Priority Effects. Annual Review of Ecology, Evolution, and Systematics. Annual Reviews Inc. pp 1–23.

Gilbert JA, Steele JA, Caporaso JG, Steinbruck L, Reeder J, Temperton B et al (2012). Defining seasonal marine microbial community dynamics. ISME J 6: 298–308.

Graham EB, Crump AR, Resch CT, Fansler S, Arntzen E, Kennedy DW et al (2016). Coupling Spatiotemporal Community Assembly Processes to Changes in Microbial Metabolism. Frontiers in Microbiology 7.

Guiot J, Cramer W (2016). Climate change: The 2015 Paris Agreement thresholds and Mediterranean basin ecosystems. Science 354: 465–468.

Hanson CA, Fuhrman JA, Horner-Devine MC, Martiny JBH (2012). Beyond biogeographic patterns: Processes shaping the microbial landscape. Nature Reviews Microbiology 10: 497–506.

Hao YQ, Zhao XF, Zhang DY (2016). Field experimental evidence that stochastic processes predominate in the initial assembly of bacterial communities. Environmental Microbiology 18: 1730–1739.

Hubbell SP (2001). The Unified Neutral Theory of Biodiversity and Biogeography. Monographs in Population Biology 32. Princeton University Press, Princeton NJ. 448 pp.

Kefi S, Rietkerk M, Alados CL, Pueyo Y, Papanastasis VP, ElAich A et al (2007). Spatial vegetation patterns and imminent desertification in Mediterranean arid ecosystems. Nature 449: 213–217.

King AJ, Freeman KR, McCormick KF, Lynch RC, Lozupone C, Knight R et al (2010). Biogeography and habitat modelling of high-alpine bacteria. Nature communications 1: 53.

Langenheder S, Szekely AJ (2011). Species sorting and neutral processes are both important during the initial assembly of bacterial communities. ISME J 5: 1086–1094.

Lee SH, Sorensen JW, Grady KL, Tobin TC, Shade A (2017). Divergent extremes but convergent recovery of bacterial and archaeal soil communities to an ongoing subterranean coal mine fire. ISME Journal 11: 1447–1459.

Leibold MA, Holyoak M, Mouquet N, Amarasekare P, Chase JM, Hoopes MF et al (2004). The metacommunity concept: a framework for multi-scale community ecology. Ecology Letters 7: 601–613.

Ma L, Guo C, Lü X, Yuan S, Wang R (2015). Soil moisture and land use are major determinants of soil microbial community composition and biomass at a regional scale in northeastern China. Biogeosciences 12: 2585–2596.

Ma X, Zhao C, Gao Y, Liu B, Wang T, Yuan T et al (2017). Divergent taxonomic and functional responses of microbial communities to field simulation of aeolian soil erosion and deposition. Molecular Ecology: n/a-n/a.

Maestre FT, Delgado-Baquerizo M, Jeffries TC, Eldridge DJ, Ochoa V, Gozalo B et al (2015). Increasing aridity reduces soil microbial diversity and abundance in global drylands. Proc Natl Acad Sci USA 112: 15684–15689.

McMurdie PJ, Holmes S (2013). phyloseq: An R Package for Reproducible Interactive Analysis and Graphics of Microbiome Census Data. PLoS ONE 8: e61217.

Moraetis D, Paranychianakis N, Nikolaidis N, Banwart S, Rousseva S, Kercheva M et al (2015). Sediment provenance, soil development, and carbon content in fluvial and manmade terraces at Koiliaris River Critical Zone Observatory. J Soils Sed 15: 347–364.

Morrison-Whittle P, Goddard MR (2015). Quantifying the relative roles of selective and neutral processes in defining eukaryotic microbial communities. ISME J 9: 2003–2011.

Nemergut DR, Schmidt SK, Fukami T, O’Neill SP, Bilinski TM, Stanish LF et al (2013). Patterns and processes of microbial community assembly. Microbiol Mol Biol Rev 77: 342–356.

Nerantzaki SD, Giannakis GV, Efstathiou D, Nikolaidis NP, Sibetheros IA, Karatzas GP et al (2015). Modeling suspended sediment transport and assessing the impacts of climate change in a karstic Mediterranean watershed. Sci Total Environ 538: 288–297.

Ofiţeru ID, Lunn M, Curtis TP, Wells GF, Criddle CS, Francis CA et al (2010). Combined niche and neutral effects in a microbial wastewater treatment community. Proc Natl Acad Sci USA 107: 15345–15350.

Oksanen J, Blanchet FG, Kindt R, Legendre P, Minchin PR, O’Hara RB et al (2013). vegan: Community Ecology Package. R package version 2.0-9. http://CRAN.R-project.org/package=vegan.

Peres-Neto PR, Legendre P, Dray S, Borcard D (2006). Variation partitioning of species data matrices: estimation and comparison of fractions. Ecology 87: 2614–2625.

Placella SA, Brodie EL, Firestone MK (2012). Rainfall-induced carbon dioxide pulses result from sequential resuscitation of phylogenetically clustered microbial groups. Proc Natl Acad Sci USA 109: 10931–10936.

Polymenakou PN, Mandalakis M, Stephanou EG, Tselepides A (2008). Particle Size Distribution of Airborne Microorganisms and Pathogens during an Intense African Dust Event in the Eastern Mediterranean. Environ Health Perspect 116: 292–296.

Price MN, Dehal PS, Arkin AP (2010). FastTree 2 ? Approximately Maximum-Likelihood Trees for Large Alignments. PLoS ONE 5: e9490.

Reddy TB, Thomas AD, Stamatis D, Bertsch J, Isbandi M, Jansson J et al (2015). The Genomes OnLine Database (GOLD) v.5: a metadata management system based on a four level (meta)genome project classification. Nucleic Acids Res 43: D1099–1106.

Roguet A, Laigle GS, Therial C, Bressy A, Soulignac F, Catherine A et al (2015). Neutral community model explains the bacterial community assembly in freshwater lakes. FEMS Microbiol Ecol 91: fiv125-fiv125.

Rousk J, Bååth E, Brookes PC, Lauber CL, Lozupone C, Caporaso JG et al (2010). Soil bacterial and fungal communities across a pH gradient in an arable soil. The ISME journal 4: 1340.

Sloan WT, Woodcock S, Lunn M, Head IM, Curtis TP (2007). Modeling taxa-abundance distributions in microbial communities using environmental sequence data. Microb Ecol 53: 443–455.

Smith TW, Lundholm JT (2010). Variation partitioning as a tool to distinguish between niche and neutral processes. ECOGRAPHY 33(4): 648–655

Stegen JC, Lin X, Konopka AE, Fredrickson JK (2012). Stochastic and deterministic assembly processes in subsurface microbial communities. ISME J 6: 1653–1664.

Stegen JC, Lin X, Fredrickson JK, Chen X, Kennedy DW, Murray CJ et al (2013). Quantifying community assembly processes and identifying features that impose them. ISME J 7: 2069–2079.

Stegen JC, Lin X, Fredrickson JK, Konopka AE (2015). Estimating and mapping ecological processes influencing microbial community assembly. Front Microbiol 6: 370.

Stegen JC, Fredrickson JK, Wilkins MJ, Konopka AE, Nelson WC, Arntzen EV et al (2016). Groundwater-surface water mixing shifts ecological assembly processes and stimulates organic carbon turnover. Nat Commun 7.

Tsiknia M, Paranychianakis NV, Varouchakis EA, Moraetis D, Nikolaidis NP (2014). Environmental drivers of soil microbial community distribution at the Koiliaris Critical Zone Observatory. FEMS Microbiology Ecology 90: 139–152.

Tsiknia M, Paranychianakis NV, Varouchakis EA, Nikolaidis NP (2015). Environmental drivers of the distribution of nitrogen functional genes at a watershed scale. FEMS Microbiol Ecol 91.

Vellend M, Srivastava DS, Anderson KM, Brown CD, Jankowski JE, Kleynhans EJ et al (2014). Assessing the relative importance of neutral stochasticity in ecological communities. Oikos 123: 1420–1430.

Wallenstein MD (2017). Managing and manipulating the rhizosphere microbiome for plant health: A systems approach. Rhizosphere 3: 230–232.

Wobbrock JO, Findlater L, Gergle D, Higgins JJ: The aligned rank transform for nonparametric factorial analyses using only anova procedures. Proceedings of the SIGCHI Conference on Human Factors in Computing Systems; Vancouver, BC, Canada. ACM: 2011.

Wubs ERJ, Van Der Putten WH, Bosch M, Bezemer TM (2016). Soil inoculation steers restoration of terrestrial Ecosystems. Nature Plants 2.

Zhou J, Liu W, Deng Y, Jiang Y-H, Xue K, He Z et al (2013). Stochastic Assembly Leads to Alternative Communities with Distinct Functions in a Bioreactor Microbial Community. mBio 4.

Zhou J, Ning D (2017). Stochastic Community Assembly: Does It Matter in Microbial Ecology? Microbiology and Molecular Biology Reviews 81.

