## Supplementary Figures and Tables for "Neutral processes and high inter-annual turnover shape the assembly of soil bacterial communities in a Mediterranean watershed"

**of**

**by**

Myrto Tsiknia*, Stilianos Fodelianakis, Nikolaos P. Nikolaidis, Nikolaos, V. Paranychianakis*

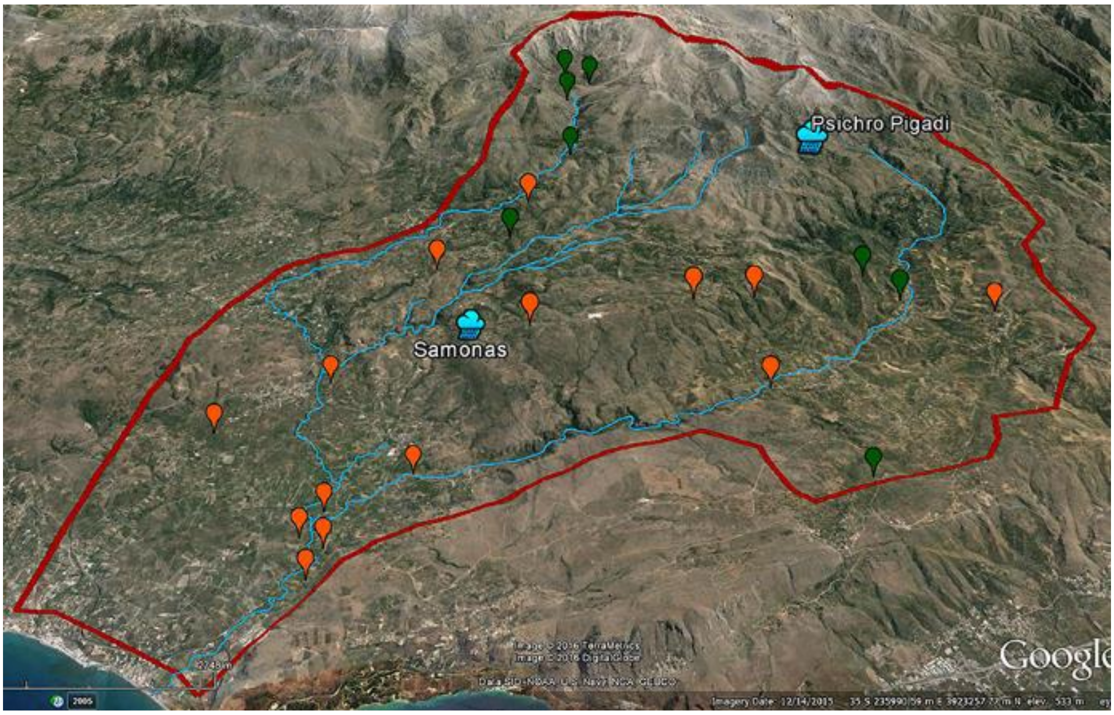


**Supplementary Figure S1**. Sampling points across Koiliaris CZO. The red-colored line indicates the borders of the watershed, while the blue one indicates the hydrographic network, the length of which approaches 36 km. Sampling points are grouped into two broad land uses, agricultural lands (orange) and natural (green) ecosystems. The two meteorological stations of the watershed are also marked at the map as raining clouds.

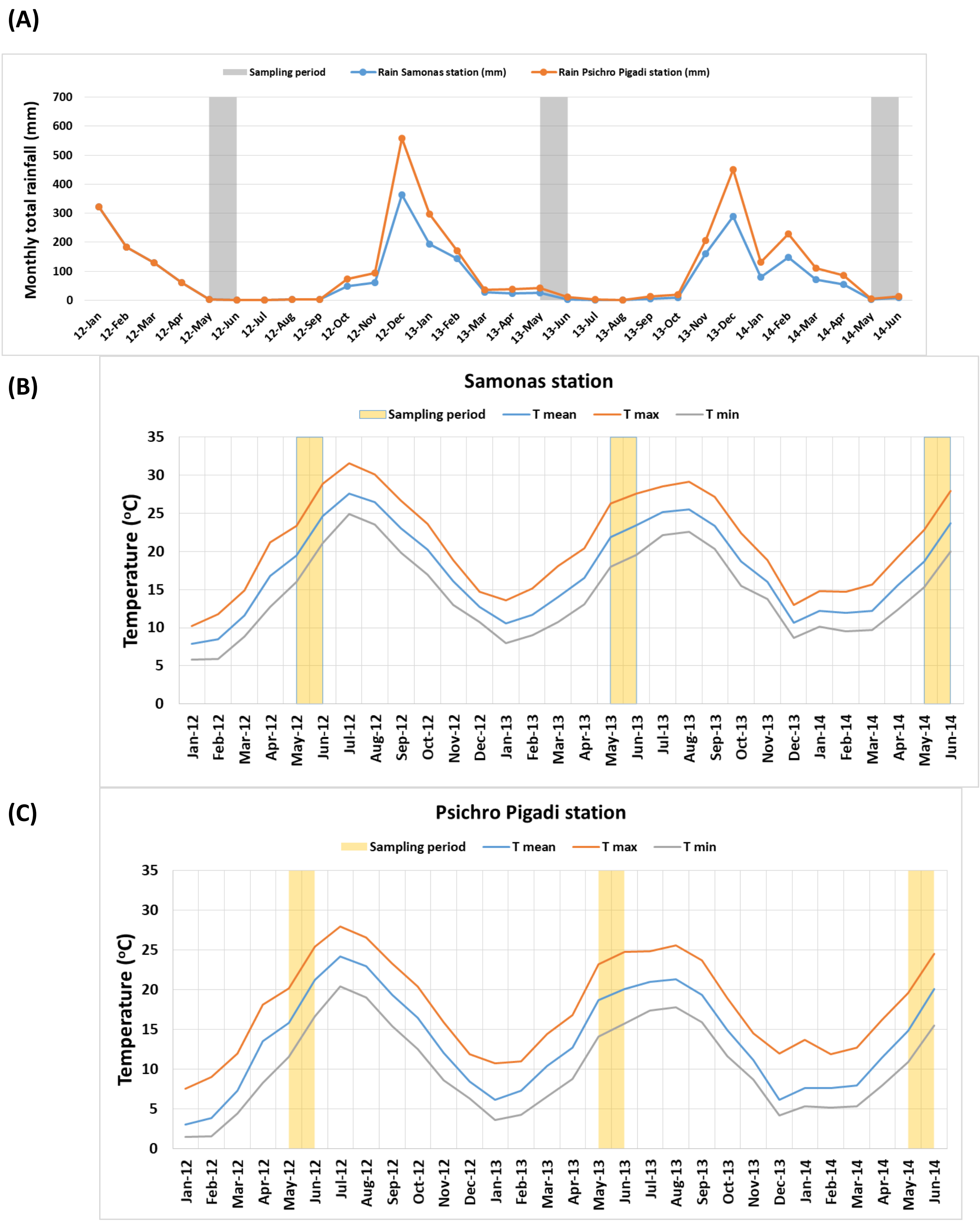


**Supplementary Figure S2**. Variation of climatic parameters in Koiliaris CZO during the period of study (2012-2014). (A) Monthly precipitation at the Samonas (blue line) and Psichro Pigadi (orange line) climatic stations. (B) Monthly temperature at Samonas and (C) Psichro Pigadi climatic stations. The location of the stations is shown in Supplementary Figure S1.

**Supplementary Table S1**. Summary statistics for the measured soil variables

| | min | max | mean±sd | CV (%) | Type of transformation |
|---|---|---|---|---|---|
| Soil moisture (%) | 1.21 | 35.9 | 9.3±4.85 | 52.7 | Log |
| pH | 5 | 8.32 | 7±0.75 | 10.7 | None |
| Electrical conductivity (EC) (μS/cm) | 36.7 | 467 | 177.1±102.7 | 57.8 | Box-Cox |
| $NO_3^-$-N (mg/kg) | 1.88 | 141 | 31±28.7 | 92.4 | Log |
| $NH_4^+$ -N (mg/kg) | 0.63 | 53.6 | 11.8±6.8 | 58.1 | Box-Cox |
| TOC (%) | 0.59 | 5.54 | 2.3±1.05 | 44.7 | Log |
| TN (%) | 0.05 | 0.4 | 0.19±0.08 | 41.67 | Box-Cox |
| C:N | 6.52 | 24.09 | 12.71±2.87 | 22.55 | Log |
| clay (%) | 9.66 | 60.55 | 34.45±13.1 | 38 | Box-Cox |
| sand (%) | 4.32 | 59.18 | 30±14.6 | 48.6 | Box-Cox |
| silt (%) | 16.11 | 90.34 | 36±9.6 | 26.7 | Log |
| Bulk density ($g/cm^3$) | 1093 | 1562 | 1328±136.2 | 10.3 | None |
| Net Nitrogen mineralization rate (NMR) (mg N/kg*d) | -3.46 | 3.05 | 0.23±0.89 | 387.88 | None |

min: minimum value; max: maximum value; CV: coefficient of variation.

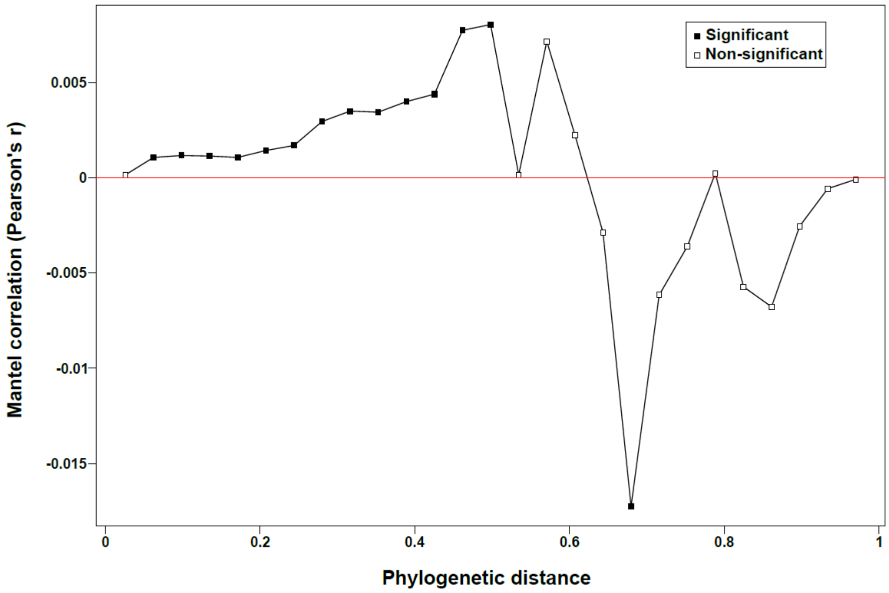


**Supplementary Figure S3**. Phylogenetic Mantel correlogram. Solid and open rectangles indicate significant and nonsignificant correlations, respectively, among OTUs niche preferences and OTUs phylogenetic distances at increasing distance classes.

### Taxonomic information

(A)

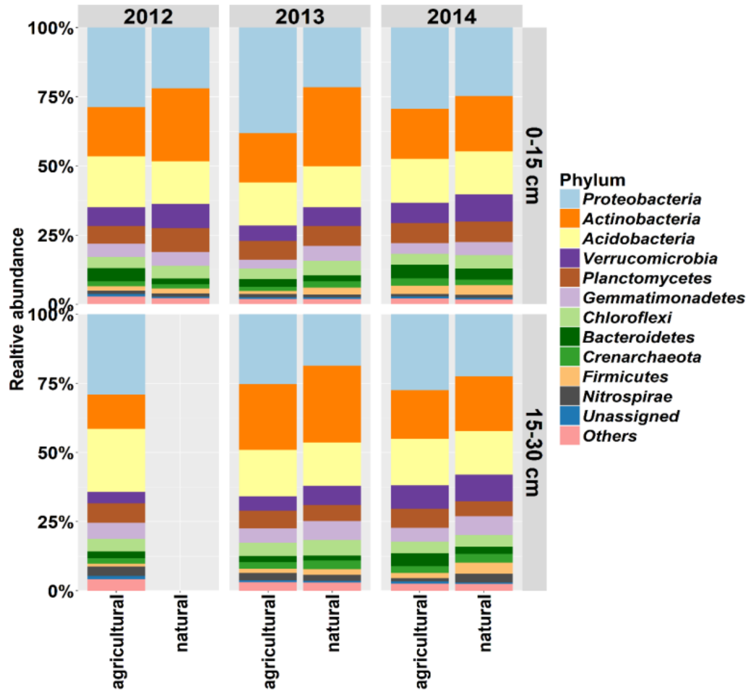


(B)

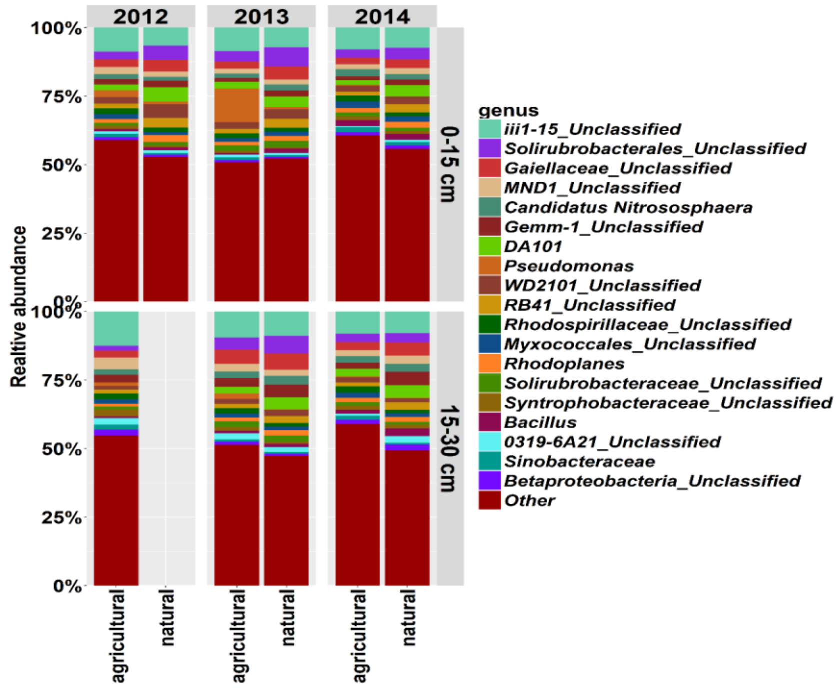


**Supplementary Figure S4**. Relative abundance of the most abundant (A) phyla (>1%) and (B) genera at Koiliaris watershed, for all the sampling years (2012, 2013, 2014), soil depths (0-15 and 15-30 cm), and land uses (agricultural vs natural).

The majority (99.3%) of reads were assigned to 48 phyla, 11 of which represented 96.8% of the total community (Supplementary Figure S4A). In detail, Proteobacteria (27.4%), Actinobacteria (20.7%) and Acidobacteria (16.4%), dominated the microbial communities, followed by Verrucomicrobia (7%), Planctomycetes (6.8%), Gemmatimonadetes (4.9%), Chloroflexi (4.4%), Bacteroidetes (3.4%), Crenarchaeota (2.3%), Firmicutes (2.1%), and Nitrospirae (1.6%). At the genus level, reads were assigned to 860 genera, 22 of which (>1%) represented 49% of the total community (Supplementary Figure S4B). Two genera affiliated to Acidobacteria and three affiliated to Actinobacteria were the most abundant, representing 10.34% and 9.2%, respectively of the whole dataset (Supplementary Figure S3B). The detailed taxonomic profile of the samples is given in Supplementary Tables S2-S6.

**Supplementary Table S2.** Mixed-effects ANOVA of α-diversity indices and soil variables.

| | Richness estimators | | Diversity estimators | | Evenness estimator | PD estimator | Soil variables | | | | | | | |
|---|---|---|---|---|---|---|---|---|---|---|---|---|---|---|
| Factors | Observed | Chao1 | Shannon | Inv_Simpson | Pielou's J | Faith's (PD) | Moisture (%) | pH | EC (µS/cm) | $NO_3$-N (mg/kg) | $NH_4^+$-N (mg/kg) | TOC (%) | TN (%) | C:N |
| Depth | ns | ns | ns | ns | ns | ns | ns | F: 7.35 ** | ns | F: 9.24 ** | ns | F: 8.45 ** | ns | F: 7.95 ** |
| Year | F: 8.31 *** | F: 12.23 *** | ns | F: 5.73 ** | F: 30.64 *** | ns | ns | ns | F: 7.72 *** | F: 15.83 *** | F: 6.81 ** | ns | F: 10.22 *** | F: 8.22 ** |
| Land | ns | ns | ns | ns | ns | ns | F: 4.72 * | ns | ns | ns | ns | F: 6.75 * | ns | F: 5.75 * |
| Depth:Year | ns | ns | ns | ns | ns | ns | ns | ns | ns | ns | F: 3.47 * | ns | ns | |
| Depth:Land | ns | ns | ns | ns | ns | ns | ns | ns | ns | F: 6.97 ** | ns | ns | ns | |
| Year:Land | ns | ns | ns | ns | ns | ns | ns | ns | ns | F: 5.51 ** | ns | ns | ns | |
| Depth: Year:Land | ns | ns | ns | ns | ns | ns | ns | ns | ns | ns | ns | ns | ns | |

Significance: ns: not significant;  *: $p<0.05$;  **: $p<0.01$;  ***: $p<0.001$

**Supplementary Table S3.** Averages and standard errors of α-diversity metrics and soil variables over years, soil depths and land uses.

| | Year | | | Land use | | Soil depth | |
|---|---|---|---|---|---|---|---|
| | 2012 | 2013 | 2014 | Agricultural | Natural ecosystems | 0-15 cm | 15-30 cm |
| α-diversity | | | | | | | |
| Observed | **2250 (±244)** | **2217 (±173)** | **1525 (±51)** | 1977 (±119) | 1912 (±147) | 1878 (±125) | 2069 (±141) |
| Chao1 | **2875 (±312)** | **2919 (±217)** | **1774 (±54)** | 2481 (±154) | 2440 (±195) | 2421 (±167) | 2536 (±177) |
| Shannon | 6.24 (±0.1) | 6.03 (±0.17) | 6.38 (±0.05) | 6.22 (±0.1) | 6.18 (±0.06) | 6.14 (±0.12) | 6.3 (±0.06) |
| Inv Simpson | **188 (±18)** | **180 (±12)** | **241 (±13)** | 218 (±10) | 278 (±12) | 207 (±11) | 204 (±12) |
| Pielou's J | **0.83 (±0.01)** | **0.81 (±0.01)** | **0.87 (±0.01)** | 0.84 (±0.008) | 0.83 (±0.007) | 0.84 (±0.009) | 0.84 (±0.006) |
| Faith's (PD) | 72.8 (±3.6) | 70.1 (±3) | 76.3 (±1.3) | 74.6 (±2) | 69.7 (±1.9) | 72.1 (±2.3) | 74.5 (±1.7) |
| Soil variables | | | | | | | |
| Soil moisture | 9.83 (±0.8) | 9.98 (±0.6) | 8.45 (±0.8) | **8.05 (±0.4)** | **12.2 (±1)** | 9.01 (±0.5) | 9.86 (±0.7) |
| pH | 7.04 (±0.2) | 7.04 (±0.1) | 7.02 (±0.1) | 7.04 (±0.08) | 6.99 (±0.1) | **6.95 (±0.09)** | **7.14 (±0.1)** |
| EC | **116.98 (±14)** | **209.72 (±16)** | **210.77 (±17)** | 203.68 (±13) | 162.27 (±16) | 181.41 (±14) | 203.71 (±15) |
| $NO_3$-N | **66.36 (±8)** | **20.55 (±2.2)** | **22.76 (±2.1)** | 33.36 (±3.6) | 26.03 (±3.2) | **35.32 (±4)** | **24.96 (±2.8)** |
| $NH_4^+$-N | **12.79 (±2)** | **9.82 (±0.8)** | **13.22 (±0.8)** | 10.96 (±0.83) | 13.49 (±0.87) | 11.87 (±0.9) | 11.60 (±0.78) |
| TOC | 2.38 (±0.2) | 2.06 (±0.14) | 2.53 (±0.15) | **1.98 (±0.09)** | **3.02 (±0.18)** | **2.52 (±0.13)** | **2.00 (±0.13)** |
| TN | **0.17 (±0.01)** | **0.16 (±0.01)** | **0.22 (±0.01)** | 0.17 (±0.01) | 0.22 (±0.01) | 0.19 (±0.01) | 0.17 (±0.01) |
| C:N | **14.35 (±0.7)** | **13.0 (±0.34)** | **11.48 (±0.4)** | **12.06 (±0.31)** | **14.11 (±0.44)** | **13.19 (±0.38)** | **12.01 (±0.3)** |
| NMR | 0.69 (±0.14) | -0.17(±0.15) | 0.4 (±0.1) | 0.26 (±0.09) | 0.18 (±0.18) | 0.31(±0.21) | 0.11 (±0.1) |

Units: Soil moisture (%). EC (μS/cm); $NO_3$-N (mg/kg); $NH_4^+$-N (mg/kg); TOC (%); TN (%); NMR (mg N/kg*d)

**Supplementary Table S4.** Relationships between α-diversity metrics and environmental parameters revealed by Spearman correlation coefficient**.**

| | Richness estimators | | | | | | Diversity estimators | | | | | |
|---|---|---|---|---|---|---|---|---|---|---|---|---|
| | 2012 | | 2013 | | 2014 | | 2012 | | 2013 | | 2014 | |
| | Observed | Chao1 | Observed | Chao1 | Observed | Chao1 | Shannon | Inv Simpson | Shannon | Inv Simpson | Shannon | Inv Simpson |
| Z | ns | ns | ns | ns | ns | ns | -0.43* | -0.44* | -0.51*** | -0.42** | ns | ns |
| Soil moisture | ns | ns | ns | ns | -0.4** | -0.35 | ns | ns | -0.35* | ns | -0.44** | -032* |
| pH | ns | ns | 0.4** | 0.35* | ns | ns | ns | ns | 0.3* | 0.32* | -0.31* | ns |
| $NO_3$-N | ns | ns | ns | ns | ns | ns | ns | ns | ns | ns | ns | ns |
| $NH_4^+$-N | ns | ns | ns | ns | -0.42** | -0.39* | ns | ns | ns | ns | ns | ns |
| TOC | ns | ns | ns | ns | -0.41** | -0.39* | ns | ns | -0.34* | ns | -0.31* | ns |
| C:N | ns | ns | ns | ns | -0.36* | -0.32* | ns | ns | ns | ns | -0.38* | -0.39** |
| Clay | -0.42* | -0.45* | ns | ns | ns | ns | -0.50* | ns | -0.33* | ns | -0.54*** | -0.51*** |
| Silt | ns | ns | ns | ns | ns | ns | ns | ns | ns | ns | ns | ns |
| Bulk density | ns | ns | ns | ns | ns | ns | ns | ns | ns | ns | ns | ns |
| NMR | ns | ns | ns | ns | ns | ns | ns | ns | ns | ns | ns | ns |

Significance: ns: not significant; *: p<0,05; **: p<0,01; ***: p<0,001
Units: Soil moisture (%); EC (μS/cm); $NO_3$-N (mg/kg); $NH_4^+$-N (mg/kg); TOC (%); clay (%); silt (%); Bulk density ($g/cm^3$); NMR (mg N/kg*d); clay and silt (%)

**Supplementary Table S4** *(continued)***.** Relationships between α-diversity metrics and environmental parameters revealed by Spearman correlation coefficient.

| | Evenness estimator (Pielou's J) | | | Phylogentic Diversity estimator (Faith's PD) | | |
|---|---|---|---|---|---|---|
| | **2012** | **2013** | 2014 | 2012 | 2013 | 2014 |
| Z | ns | ns | -0.3* | -0.51* | -0.55*** | -0.33* |
| Soil moisture | ns | ns | ns | ns | -0.41*** | -0.51*** |
| pH | ns | ns | ns | ns | ns | ns |
| $NO_3$-N | ns | ns | ns | ns | ns | ns |
| $NH_4^+$-N | ns | ns | ns | ns | ns | -0.42** |
| TOC | ns | ns | ns | ns | -0.48*** | -0.47** |
| C:N | ns | ns | ns | ns | ns | -0.44** |
| Clay | -0.42* | ns | -0.6*** | -0.54** | -0.41*** | -0.57*** |
| Silt | ns | ns | ns | ns | ns | ns |
| Bulk density | ns | ns | ns | ns | ns | ns |
| NMR | ns | ns | ns | ns | ns | ns |

Significance: ns: not significant; *: *p<0.05*; **: *p<0.01*; ***: *p<0.001*
Units: Soil moisture (%); EC (µS/cm); $NO_3$-N (mg/kg); $NH_4^+$-N (mg/kg); TOC (%); silt (%); Bulk density ($g/cm^3$); NMR (mg N/kg*d); clay and silt (%)

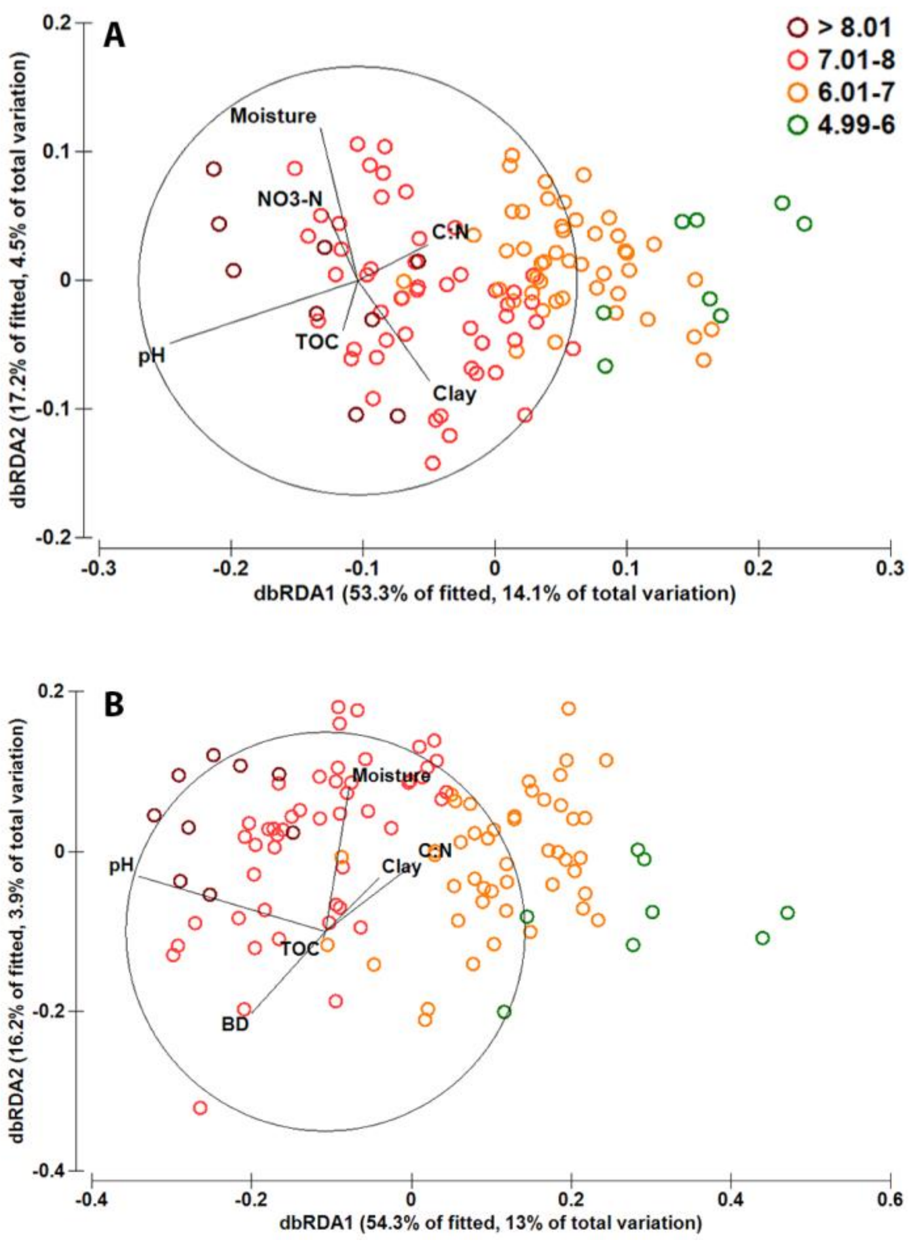


**Supplementary Figure S5**. RDA plots of A) weighted UniFrac distance and B) Bray-Curtis dissimilarities of microbial community in Koiliaris CZO. Each point represents a different sample. Samples have been grouped according to pH range. Only the significant variables have been included in the plots.

**Supplementary Table S5.** PERMANOVA (999 permutations) of the effect of sampling year, soil depth and land use on the UniFrac distances and Bray-Curtis dissimilarities**.**

| | Weighted UniFrac | Bray-Curtis |
|---|---|---|
| Year | 5.7% *** | 10% *** |
| Land use | 4.7% *** | 3% *** |
| Soil depth | 2.6% ** | 1.8% ** |
| Significance: **: *p<0.01*; ***: *p<0.001* |  |  |

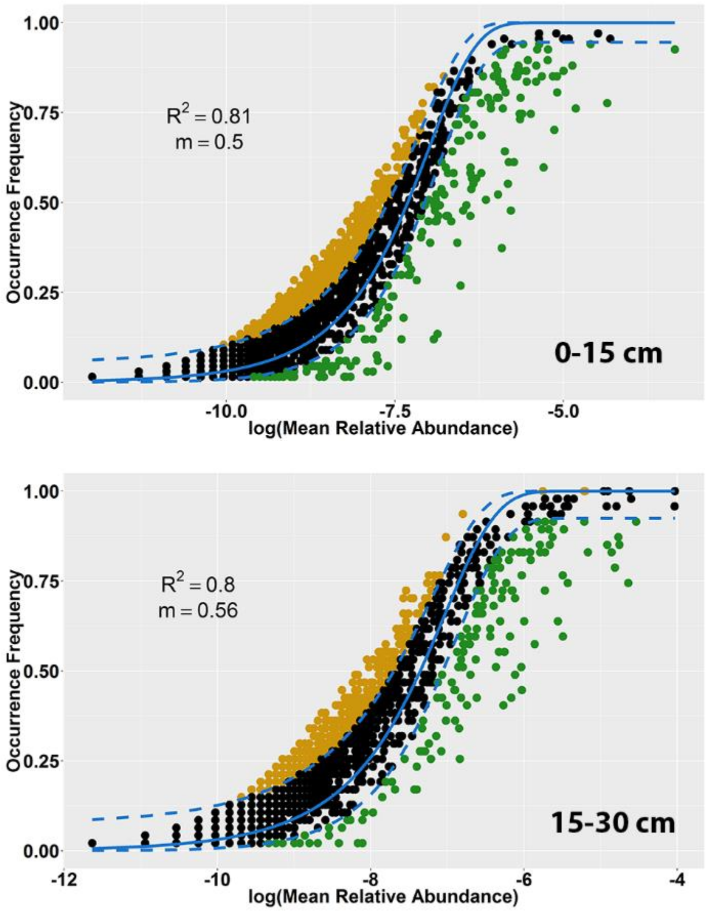


**Supplementary Figure S6**. Fit of Sloan Neutral Community Model for Prokaryotes for each soil depth. In each plot model's fit ($R^2$) and migration rate (m) are shown. OTUs that occur more frequently than predicted by the model are shown in yellow, while those that found to occur less frequently than predicted are shown in green. Dashed blue lines represent the 95% confidence intervals of the model prediction (solid blue line).

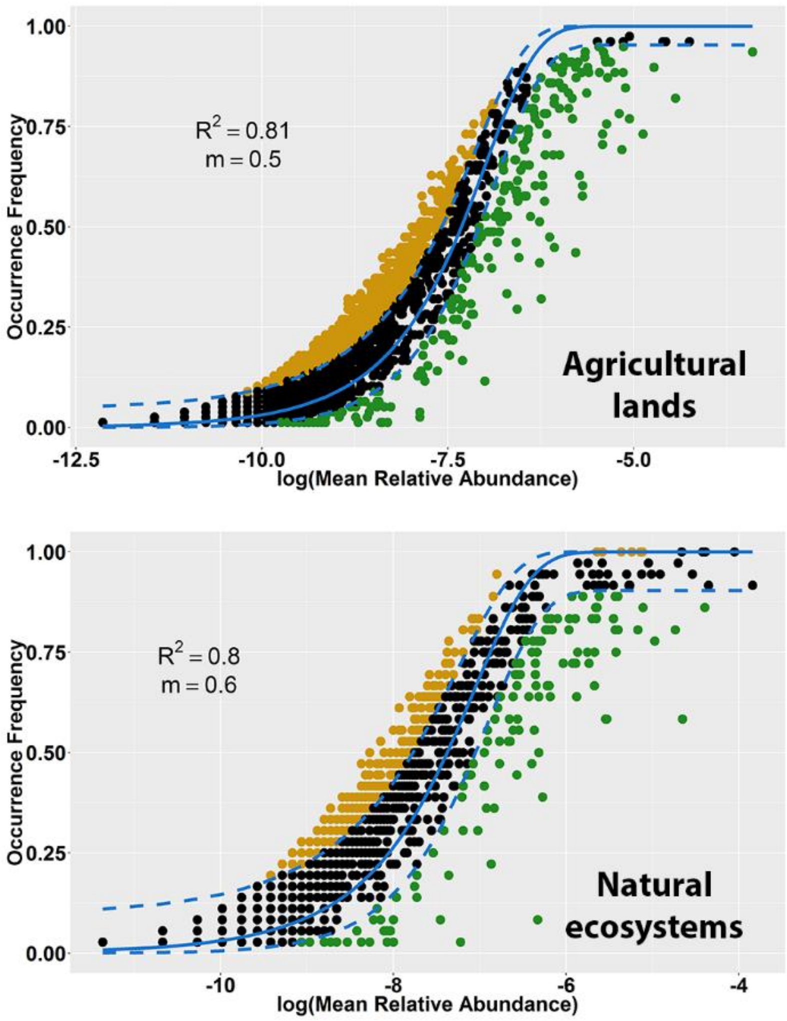


**Supplementary Figure S7**. Fit of Sloan Neutral Community Model for Prokaryotes for land use. In each plot model's fit ($R^2$) and migration rate (m) are shown. OTUs that occur more frequently than predicted are shown in yellow while those found to occur less frequently than predicted are shown in green. Dashed blue lines represent the 95% confidence intervals of the model prediction (solid blue line).

**Supplementary Table S6**. Parameters and fits of neutral and binomial models. The estimations were performed by the code provided by Burns et al. 2016.

| | Year | | | Land Use | | Soil Depth | |
|---|---|---|---|---|---|---|---|
| SNCM parameters | 2012 | 2013 | 2014 | Agricultural land | Natural ecosystem | 0-15 cm | 15-30 cm |
| m | 0.36 | 0.38 | 0.60 | 0.50 | 0.60 | 0.49 | 0.56 |
| m.ci | 0.02 | 0.02 | 0.03 | 0.02 | 0.03 | 0.02 | 0.02 |
| m.mle | 0.36 | 0.38 | 0.60 | 0.50 | 0.60 | 0.49 | 0.56 |
| maxLL | -5375 | -5087 | -5318 | -12684 | -6577 | -11294 | -9421 |
| binoLL | -3835 | -4041 | -4933 | -10802 | -6607 | -9377 | -8741 |
| poisLL | -3835 | -4042 | -4934 | -10804 | -6609 | -9379 | -8742 |
| Rsqr | 0.678 | 0.685 | 0.801 | 0.812 | 0.797 | 0.809 | 0.804 |
| Rsqr.bino | 0.234 | 0.374 | 0.663 | 0.653 | 0.749 | 0.626 | 0.707 |
| Rsqr.pois | 0.234 | 0.374 | 0.663 | 0.653 | 0.749 | 0.626 | 0.708 |
| RMSE | 0.089 | 0.089 | 0.084 | 0.061 | 0.080 | 0.064 | 0.070 |
| RMSE.bino | 0.137 | 0.125 | 0.109 | 0.084 | 0.089 | 0.089 | 0.085 |
| RMSE.pois | 0.137 | 0.125 | 0.109 | 0.083 | 0.089 | 0.089 | 0.085 |
| AIC | -10746 | -10171 | -10632 | -25365 | -13551 | -22584 | -18838 |
| BIC | -10733 | -10157 | -10619 | -25351 | -13538 | -22570 | -18824 |
| AIC.bino | -7666 | -8078 | -9862 | -21601 | -13211 | -18751 | -17478 |
| BIC.bino | -7652 | -8065 | -9849 | -21586 | -13198 | -18736 | -17464 |
| AIC.pois | -7667 | -8080 | -9864 | -21605 | -13214 | -18754 | -17481 |
| BIC.pois | -7654 | -8067 | -9851 | -21591 | -13201 | -18740 | -17467 |
| N | 2399 | 2399 | 2399 | 2399 | 2399 | 2399 | 2399 |
| Samples | 22 | 22 | 22 | 78 | 36 | 67 | 47 |
| Richness | 5363 | 5064 | 5006 | 9258 | 5922 | 8462 | 7559 |

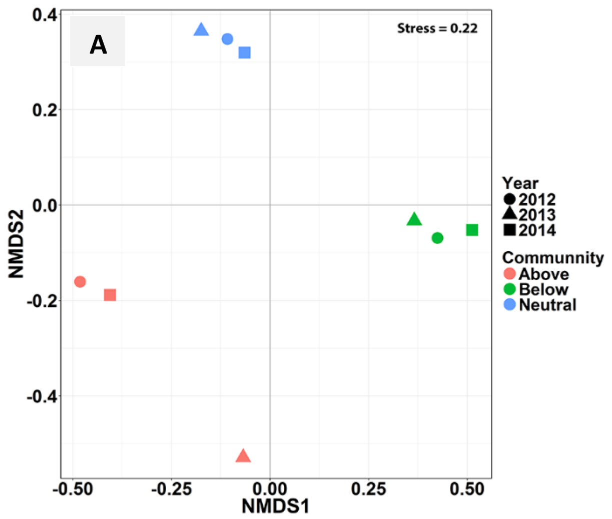


**Supplementary Figure S8**. NMDS plot of Bray-Curtis dissimilarities (A) and weighted Unifrac distance (B) of the neutral, less frequent than expected and more frequent than expected microbial communities as they were separated by the SNCM.

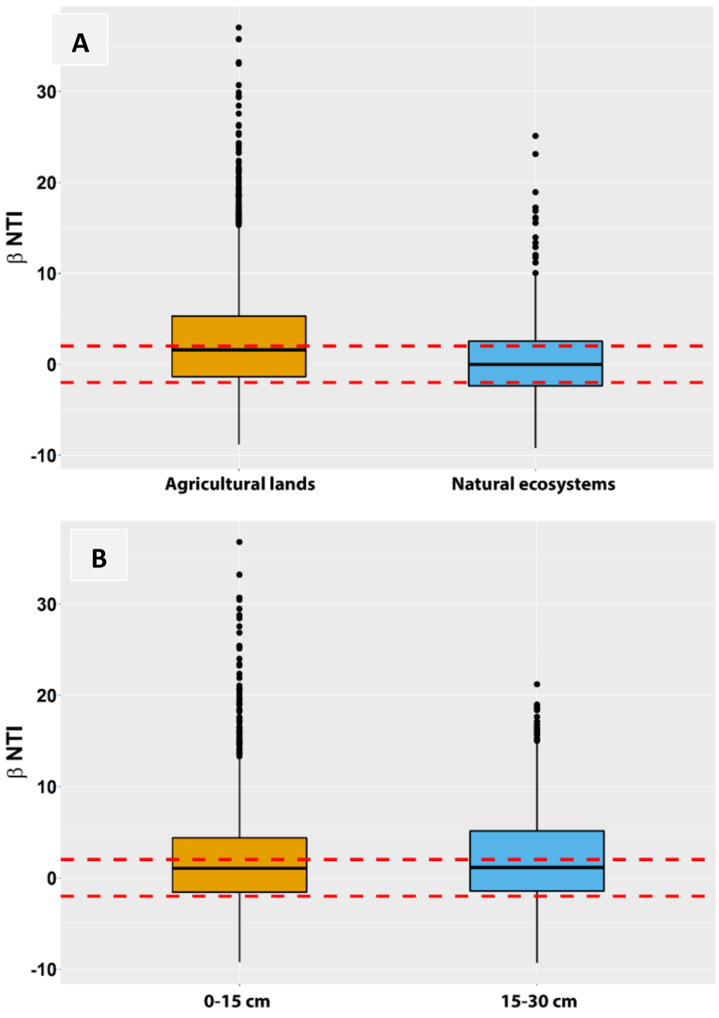


**Supplementary Figure S9**. Box plots of βNTI distributions between land uses (A) and soil depths (B) showing the median (thick black line), the first quartile (lower box bound), the third quartile (upper box bound), and outliers (black circles). Horizontal red dashed lines indicate upper and lower significance thresholds of βNTI at +2 and −2, respectively.

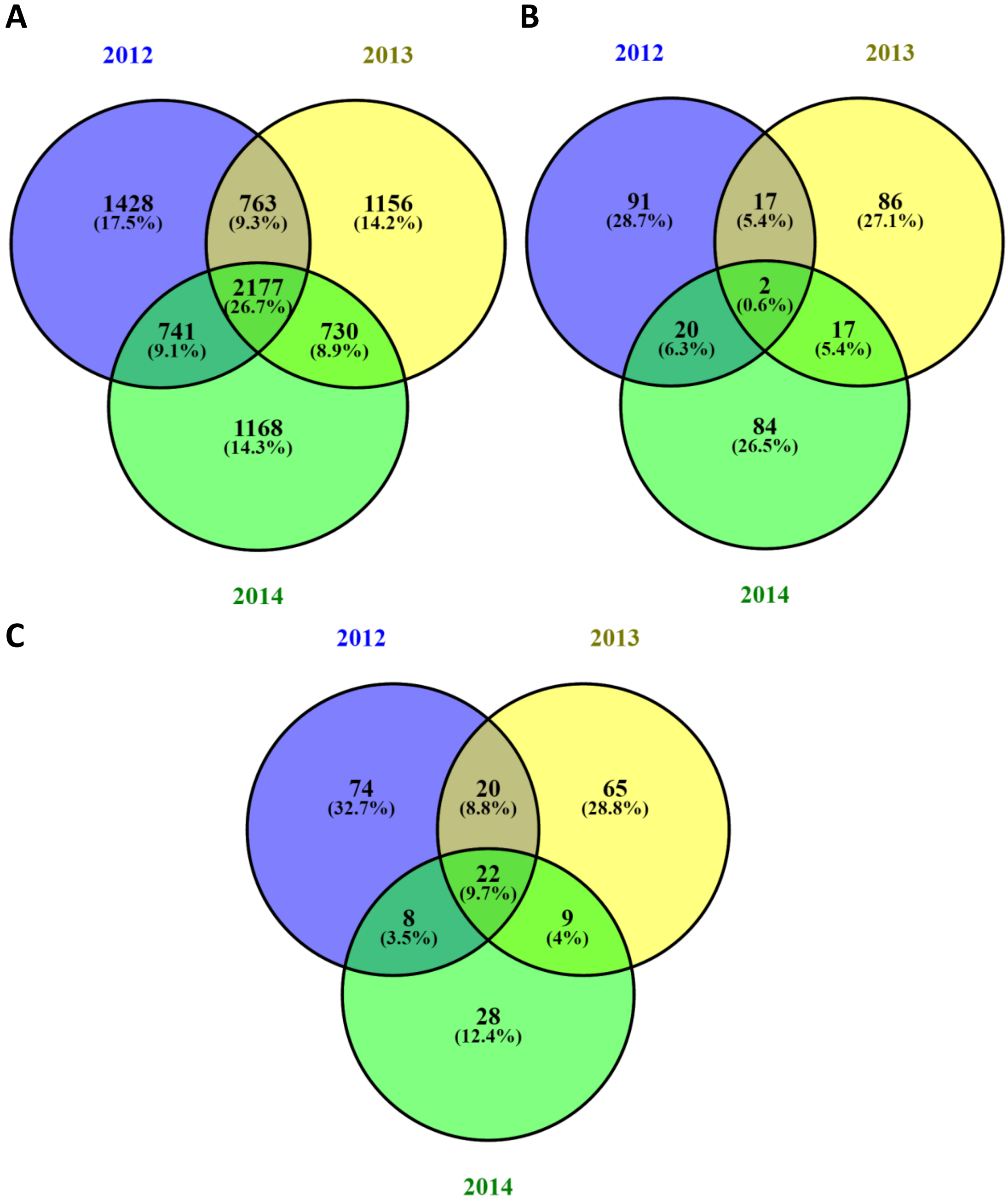


**Supplementary Figure S10**. Venn diagrams of the common OTUs for the three sampling years; (A) the neutral community, (B) the more frequent than expected OTUs and (B) the less frequent than expected OTUs. Venn diagrams were constructed with the VENNY interactive tool (v2.1) (Oliveros, 2007)

**Supplementary Table S7**. Summary of model selection results for the whole dataset, each land use and each soil depth. Forward variable selection performed with *ordiR2step()* function, with 999 permutations, separately for soil and spatial variables.

| | Soil variables | Spatial variables |
|---|---|---|
| All data | pH$^{**}$, silt$^{**}$, TOC$^{**}$, soil moisture$^{**}$, C:N$^{**}$, $NO_3^-$-N$^{*}$ | $V_1^{**}$, $V_2^{**}$, $V_3^{**}$, $V_4^{**}$, $V_5^{**}$, $V_6^{**}$, $V_7^{**}$ |
| Agricultural land | pH$^{**}$, TOC$^{**}$, C:N$^{**}$, soil moisture$^{*}$ | $V_1^{**}$, $V_2^{**}$, $V_4^{**}$, $V_5^{**}$, $V_6^{**}$, $V_7^{**}$ |
| Natural ecosystem | pH$^{**}$, TOC$^{**}$, $NO_3^-$-N$^{**}$, soil moisture$^{**}$ | $V_1^{**}$, $V_3^{**}$ |
| 0-15 cm | pH$^{**}$, TOC$^{**}$, $NO_3^-$-N$^{**}$, soil moisture$^{**}$, C:N$^{**}$ | $V_1^{**}$, $V_2^{**}$, $V_3^{*}$, $V_6^{**}$, $V_7^{*}$, |
| 15-30 cm | pH$^{**}$, clay$^{**}$, soil moisture$^{**}$, TOC$^{**}$ | $V_1^{**}$, $V_2^{**}$, $V_3^{**}$, $V_4^{*}$, $V_6^{**}$ |

Significance: ns: not significant; *: *p<0.05*; **: *p<0.01*; ***: *p<0.001*
For spatial variables $V_x$ refers to the number of the PCNM vector

**(A)**

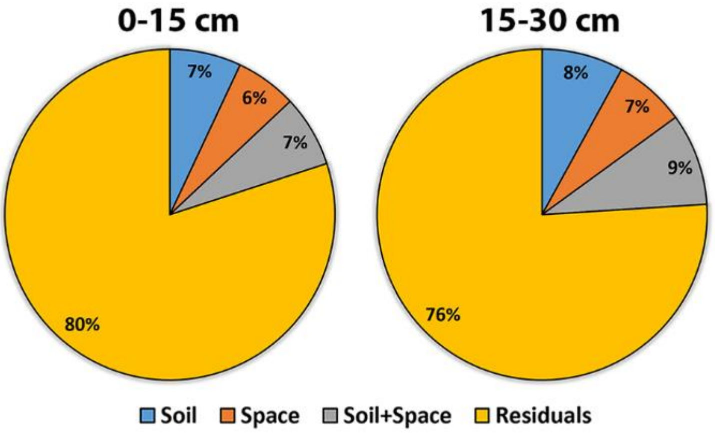


**(B)**

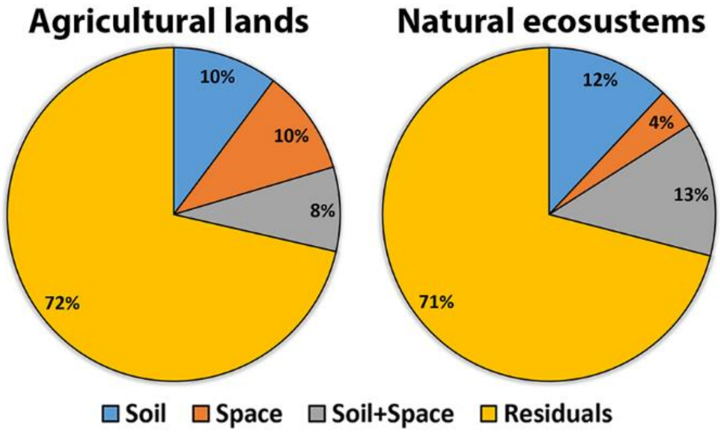


**Supplementary Figure S11**. Variation partitioning analysis of the soil depth (A) and land use (B) illustrating the effects of soil and spatial variables (space) and the shared effects on β-diversity. Values refer to the percentage of variation explained by each fraction, including the pure soil effect (Soil), the shared effect between soil and space (Soil+Space), the pure spatial effect (Space), and the unexplained variation (Residuals).

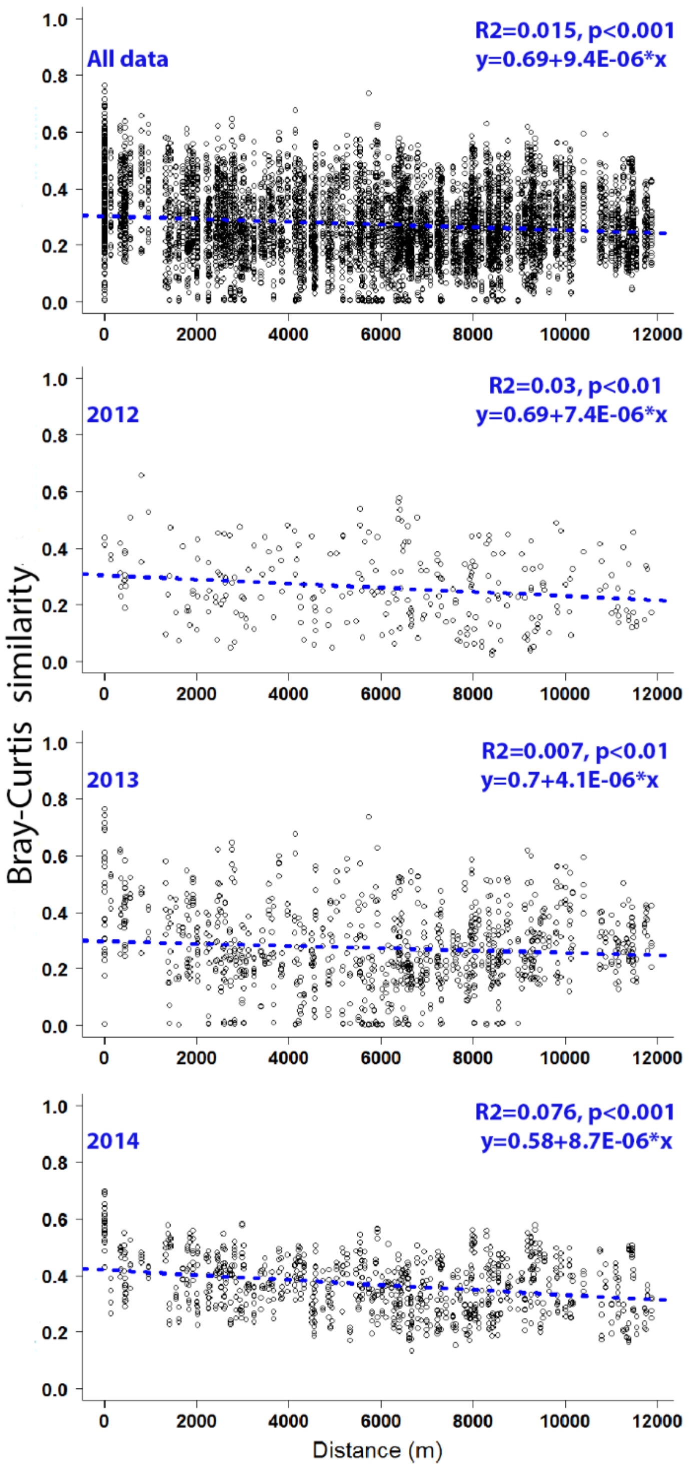


**Supplementary Figure S12**. Distance-decay plots of Bray-Curtis similarity, for the whole dataset and for each year separately. Dashed lines correspond to linear regression models. The fit and the equation of each linear model are shown on the right edge of each plot. On the equations y corresponds to Bray-Curtis similarity, while x stands for distance.

**Supplementary Table S8**. Relative contribution of individual mechanisms of assembly (dispersal, selection, drift) on soil microbial communities as they are affected by land use and soil depth. The estimation is based on the framework of Stegen et al (2015).

| | Land use | | Soil depth | |
|---|---|---|---|---|
| | Agricultural | Natural | 0-15 cm | 15-30 cm |
| Variable selection (βNTI> +2) | 47.3 | 30.6 | 42.0 | 46.4 |
| Homogenous selection (βNTI< -2) | 21.1 | 29.6 | 23.3 | 20.4 |
| **Total selection** | 68.4 | 60.2 | 65.3 | 66.8 |
| Dispersal limitation (βNTI<\|2\| & RC> +0.95) | 13.2 | 19.8 | 15.1 | 17.6 |
| Dispersal homogenizing (βNTI<\|2\| & RC< -0.95) | 7.2 | 8.9 | 6.9 | 6.4 |
| Undominated (βNTI<\|2\| & RC< \|0.95\|) | 12.3 | 11.1 | 12.7 | 9.2 |
| **Total neutral** | 31.7 | 39.8 | 34.7 | 33.2 |

the ‘undominated’ fraction being the residual fraction of comparisons after accounting for all other assembly processes

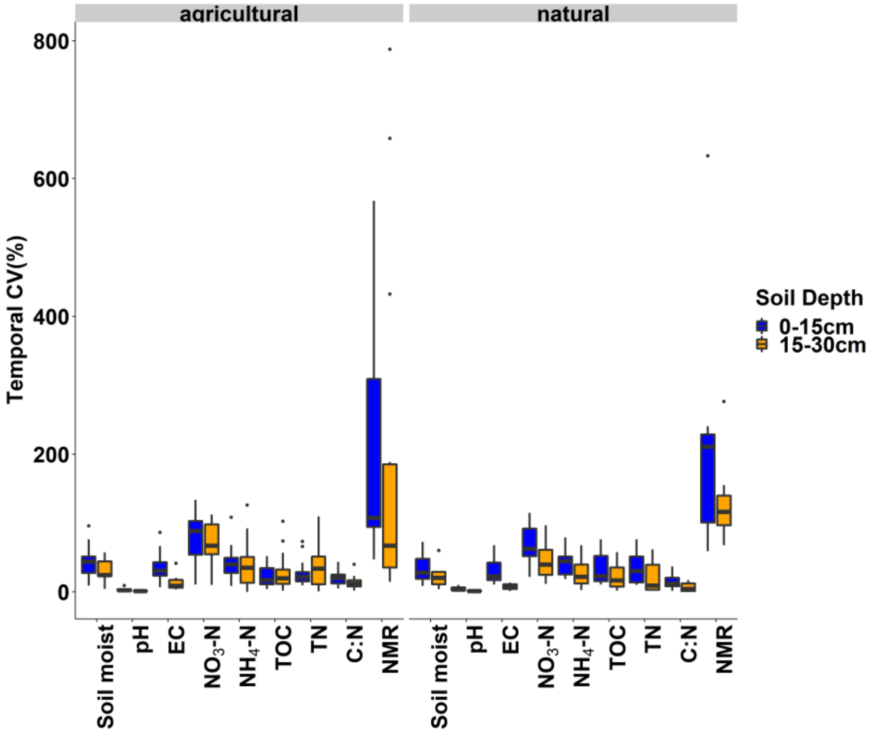


**Supplementary Figure S13** Distribution of the temporal coefficient of variation (CV %) of the measured soil variables. The temporal CV calculated from the three annual replicates of each sampling site. The upper and lower boundaries indicate the 75th and the 25th percentile; the mid-line indicates the median, while the cross indicates the mean of the distribution; above and below whiskers indicate the 90th and 10th percentiles; dots indicate values identified as outliers.

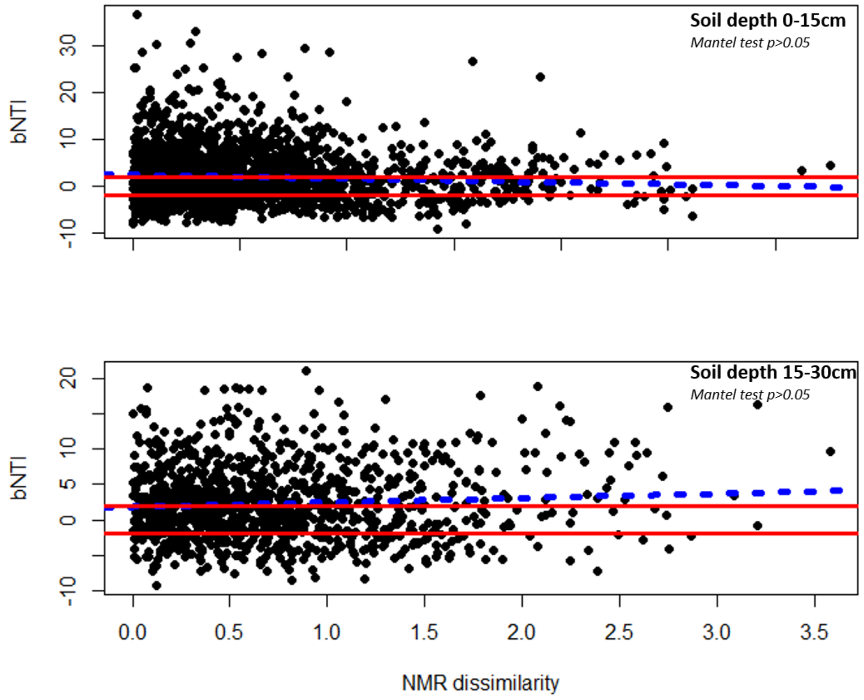


**Supplementary Figure S14** The relationships between the βNTI and NMR dissimilarity (Euclidean distance) for each soil depth. The regression slopes of the linear relationships based on Gaussian generalized model are shown with blue dashed lines (non-significant; Mantel test; 9999 permutations; p> 0.05). The solid red line indicates the upper and lower significance thresholds at βNTI =+2 and −2, respectively.

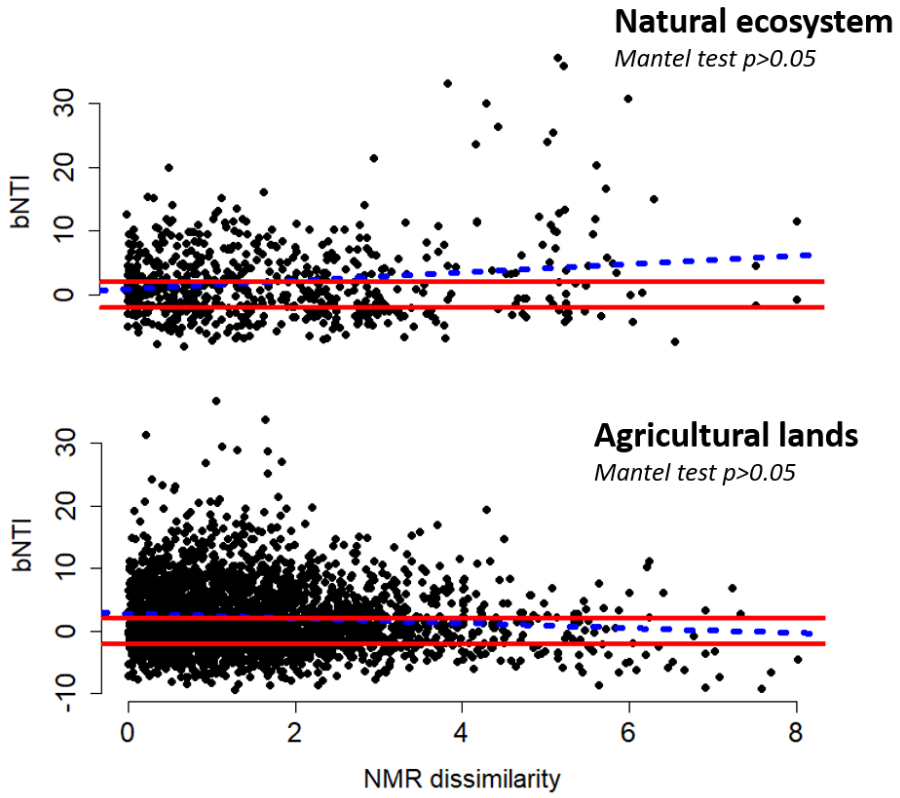

**Supplementary Figure S15** The relationships between the βNTI and NMR dissimilarity (Euclidean distance) for each land use. The regression slopes of the linear relationships based on Gaussian generalized model are shown with blue dashed lines (non-significant; Mantel test; 9999 permutations; $p > 0.05$). The solid red line indicates the highest and lowest significance thresholds at βNTI =+2 and −2, respectively.

### References

Burns, A.R., Stephens, W.Z., Stagaman, K., Wong, S., Rawls, J.F., Guillemin, K., Bohannan, B.J., 2016. Contribution of neutral processes to the assembly of gut microbial communities in the zebrafish over host development. ISME J 10, 655-664

Oliveros, J.C. Venny., 2007 An interactive tool for comparing lists with Venn's diagrams. http://bioinfogp.cnb.csic.es/tools/venny/index.html
