## Supplementary Table S15 for "Neutral processes and high inter-annual turnover shape the assembly of soil bacterial communities in a Mediterranean watershed"

**Supplementary Table S9 Significant correlations between individual taxa at a high taxonomic level (Phylum or Class) and NMR**

| | | | | |
|---|---|---|---|---|
| PHYLUM_LEVEL | | | | |
| ALL _SOIL | | | | |
| | | | *r* | *p* |
| | Actinobacteria | NMR | -0.283 | 0.002 |
| | Chloroflexi | NMR | -0.310 | 0.001 |
| | Elusimicrobia | NMR | 0.277 | 0.003 |
| AGRICULTURE _SOIL | | | | |
| | | | *r* | *p* |
| | Chloroflexi | NMR | -0.279 | 0.013 |
| | Cyanobacteria | NMR | 0.227 | 0.046 |
| | Elusimicrobia | NMR | 0.342 | 0.002 |
| | Spirochaetes | NMR | 0.224 | 0.049 |
| NATURAL _SOIL | | | | |
| | | | *r* | *p* |
| | Actinobacteria | NMR | -0.348 | 0.038 |
| | Chloroflexi | NMR | -0.337 | 0.044 |
| | Verrucomicrobia | NMR | 0.333 | 0.047 |
| 0-15CM _SOIL | | | | |
| | | | *r* | *p* |
| | Actinobacteria | NMR | -0.272 | 0.026 |
| | Chloroflexi | NMR | -0.255 | 0.037 |
| | Elusimicrobia | NMR | 0.244 | 0.047 |
| | GAL15 | NMR | -0.266 | 0.030 |
| 15-30CM _SOIL | | | | |
| | | | *r* | *p* |
| | Euryarchaeota | NMR | 0.292 | 0.047 |
| | Chloroflexi | NMR | -0.353 | 0.015 |
| | Elusimicrobia | NMR | 0.294 | 0.045 |
| 2012_depth_0-15 | | | | |
| | Acidobacteria | NMR | 0.586 | 0.004 |
| | Actinobacteria | NMR | -0.427 | 0.047 |
| | Chloroflexi | NMR | -0.429 | 0.047 |
| | WS3 | NMR | 0.470 | 0.027 |
| | [Caldithrix] | NMR | 0.497 | 0.019 |
| | [Thermi] | NMR | -0.428 | 0.047 |
| 2013_depth_0-15 | | | | |
| | GAL15 | NMR | -0.418 | 0.047 |
| 2013_depth_15-30 | | | | |
| | Euryarchaeota | NMR | 0.478 | 0.021 |

| | | | | |
|---|---|---|---|---|
| | Armatimonadetes | NMR | -0.481 | 0.020 |
| | Chloroflexi | NMR | -0.497 | 0.016 |
| 2014_depth_0-15 | | | | |
| | Cyanobacteria | NMR | 0.451 | 0.035 |
| | Fibrobacteres | NMR | 0.454 | 0.034 |
| | WS3 | NMR | -0.508 | 0.016 |
| 2014_depth_15-30 | | | | |
| | Actinobacteria | NMR | -0.466 | 0.029 |
| | Elusimicrobia | NMR | 0.435 | 0.043 |
| | Spirochaetes | NMR | 0.473 | 0.026 |
| agriculture_depth_0-15 | | | | |
| | Cyanobacteria | NMR | 0.353 | 0.017 |
| | Gemmatimonadetes | NMR | 0.369 | 0.013 |
| | NC10 | NMR | -0.323 | 0.031 |
| agriculture_depth_15-30 | | | | |
| | Elusimicrobia | NMR | 0.426 | 0.013 |
| | Spirochaetes | NMR | 0.354 | 0.043 |
| natural_depth_0-15 | | | | |
| | NONE | | | |
| natural_depth_15-30 | | | | |
| | Unclassified | NMR | -0.579 | 0.030 |

| CLASS_LEVEL | | | | |
|---|---|---|---|---|
| ALL _SOIL | | | | |
| | | | *r* | *p* |
| | Chlorobi;c__BSV26 | NMR | 0.253 | 0.007 |
| | Chloroflexi;Other | NMR | -0.333 | 0.000 |
| | Chloroflexi;c__Ellin6529 | NMR | -0.232 | 0.013 |
| | Elusimicrobia;Other | NMR | 0.268 | 0.004 |
| | Elusimicrobia;c__Elusimicrobia | NMR | 0.276 | 0.003 |
| | Proteobacteria;c__Betaproteobacteria | NMR | 0.190 | 0.042 |
| | Proteobacteria;c__Deltaproteobacteria | NMR | 0.215 | 0.022 |
| | Verrucomicrobia;c__[Pedosphaerae] | NMR | 0.212 | 0.024 |
| AGRICULTURE _SOIL | | | | |
| | | | *r* | *p* |
| | Armatimonadetes;c__[Fimbriimonadia] | NMR | 0.230 | 0.043 |
| NATURAL _SOIL | | | | |
| | | | *r* | *p* |
| | Acidobacteria;c__S035 | NMR | -0.610 | 0.000 |
| | Actinobacteria;c__Acidimicrobiia | NMR | -0.358 | 0.032 |
| 0-15CM _SOIL | | | | |
| | | | *r* | *p* |
| | Euryarchaeota;c__Methanobacteria | NMR | 0.304 | 0.012 |
| | Acidobacteria;c__ | NMR | 0.246 | 0.045 |
| | Acidobacteria;c__Sva0725 | NMR | 0.250 | 0.042 |
| | Armatimonadetes;c__[Fimbriimonadia] | NMR | 0.292 | 0.017 |
| | Bacteroidetes;c__At12OctB3 | NMR | 0.246 | 0.045 |
| 15-30CM _SOIL | | | | |
| | | | *r* | *p* |
| | Bacteroidetes;c__At12OctB3 | NMR | -0.326 | 0.025 |
| 2012_depth_0-15 | | | | |
| | Acidobacteria;c__ | NMR | 0.563 | 0.006 |
| | Acidobacteria;c__AT-s54 | NMR | 0.571 | 0.006 |
| | Acidobacteria;c__Acidobacteria-6 | NMR | 0.615 | 0.002 |
| | Acidobacteria;c__Acidobacteriia | NMR | -0.430 | 0.046 |
| | Acidobacteria;c__RB25 | NMR | 0.505 | 0.017 |
| | Acidobacteria;c__S035 | NMR | 0.544 | 0.009 |
| | Actinobacteria;c__Actinobacteria | NMR | -0.571 | 0.006 |
| | Armatimonadetes;c__0319-6E2 | NMR | 0.458 | 0.032 |
| | Bacteroidetes;Other | NMR | 0.498 | 0.018 |
| | Chlorobi;c__BSV26 | NMR | 0.465 | 0.029 |
| | Chloroflexi;c__Ktedonobacteria | NMR | -0.437 | 0.042 |
| | Chloroflexi;c__S085 | NMR | 0.532 | 0.011 |

| | | | | |
|---|---|---|---|---|
| | Chloroflexi;c__SAR202 | NMR | 0.448 | 0.036 |
| | Gemmatimonadetes;c__ | NMR | 0.597 | 0.003 |
| | Gemmatimonadetes;c__Gemm-2 | NMR | 0.546 | 0.009 |
| | Gemmatimonadetes;c__Gemm-3 | NMR | 0.461 | 0.031 |
| | Gemmatimonadetes;c__Gemm-5 | NMR | 0.619 | 0.002 |
| | Planctomycetes;c__ | NMR | 0.501 | 0.017 |
| | Planctomycetes;c__BD7-11 | NMR | 0.444 | 0.039 |
| | Planctomycetes;c__Pla3 | NMR | 0.446 | 0.038 |
| | Proteobacteria;c__ | NMR | 0.430 | 0.046 |
| | Proteobacteria;c__Deltaproteobacteria | NMR | 0.447 | 0.037 |
| | WS3;c__PRR-12 | NMR | 0.470 | 0.027 |
| | [Caldithrix];c__KSB1 | NMR | 0.497 | 0.019 |
| | [Thermi];c__Deinococci | NMR | -0.428 | 0.047 |
| 2013_depth_0-15 | | | | |
| | Bacteroidetes;c__At12OctB3 | NMR | 0.432 | 0.039 |
| | Bacteroidetes;c__VC2_1_Bac22 | NMR | -0.451 | 0.031 |
| | Chloroflexi;Other | NMR | -0.653 | 0.001 |
| | Chloroflexi;c__Gitt-GS-136 | NMR | -0.505 | 0.014 |
| | GAL15;c__ | NMR | -0.418 | 0.047 |
| 2013_depth_15-30 | | | | |
| | Euryarchaeota;c__Thermoplasmata | NMR | 0.478 | 0.021 |
| | Bacteroidetes;c__Flavobacteriia | NMR | -0.414 | 0.049 |
| | Chloroflexi;c__Ellin6529 | NMR | -0.433 | 0.039 |
| | Verrucomicrobia;c__[Methylacidiphilae] | NMR | -0.476 | 0.022 |
| 2014_depth_0-15 | | | | |
| | Acidobacteria;c__Acidobacteria-6 | NMR | -0.434 | 0.043 |
| | Acidobacteria;c__BPC102 | NMR | -0.578 | 0.005 |
| | Acidobacteria;c__RB25 | NMR | -0.492 | 0.020 |
| | Acidobacteria;c__S035 | NMR | -0.495 | 0.019 |
| | Acidobacteria;c__Sva0725 | NMR | 0.545 | 0.009 |
| | Chloroflexi;c__TK17 | NMR | -0.590 | 0.004 |
| | Fibrobacteres;c__Fibrobacteria | NMR | 0.454 | 0.034 |
| | WS3;c__PRR-12 | NMR | -0.508 | 0.016 |
| 2014_depth_15-30 | | | | |
| | Acidobacteria;c__[Chloracidobacteria] | NMR | -0.432 | 0.045 |
| | Actinobacteria;c__Rubrobacteria | NMR | -0.469 | 0.028 |
| | Actinobacteria;c__Thermoleophilia | NMR | -0.506 | 0.016 |
| | Chloroflexi;c__ | NMR | -0.573 | 0.005 |
| | Chloroflexi;c__Chloroflexi | NMR | -0.430 | 0.046 |
| | Chloroflexi;c__TK10 | NMR | -0.527 | 0.012 |
| | Elusimicrobia;c__Elusimicrobia | NMR | 0.451 | 0.035 |
| | Gemmatimonadetes;c__ | NMR | 0.442 | 0.039 |
| | OP3;c__koll11 | NMR | 0.463 | 0.030 |
| | Proteobacteria;c__Deltaproteobacteria | NMR | 0.468 | 0.028 |
| | Spirochaetes;c__[Leptospirae] | NMR | 0.500 | 0.018 |

| | | | | |
|---|---|---|---|---|
| agriculture_depth_0-15 | | | | |
| | Acidobacteria;c__ | NMR | 0.332 | 0.026 |
| | Chloroflexi;c__C0119 | NMR | 0.300 | 0.045 |
| | Gemmatimonadetes;c__Gemm-1 | NMR | 0.418 | 0.004 |
| | NC10;c__12-24 | NMR | -0.325 | 0.030 |
| agriculture_depth_15-30 | | | | |
| | Chlorobi;c__BSV26 | NMR | 0.355 | 0.043 |
| | Chloroflexi;c__C0119 | NMR | -0.374 | 0.032 |
| | Elusimicrobia;c__Elusimicrobia | NMR | 0.451 | 0.008 |
| | Planctomycetes;c__C6 | NMR | 0.454 | 0.008 |
| | Proteobacteria;c__Betaproteobacteria | NMR | 0.376 | 0.031 |
| | Proteobacteria;c__Deltaproteobacteria | NMR | 0.464 | 0.006 |
| natural_depth_0-15 | | | | |
| | Euryarchaeota;c__Methanobacteria | NMR | 0.452 | 0.035 |
| | Acidobacteria;c__S035 | NMR | -0.613 | 0.002 |
| | Acidobacteria;c__Sva0725 | NMR | 0.429 | 0.047 |
| | Armatimonadetes;c__[Fimbriimonadia] | NMR | 0.447 | 0.037 |
| | Chloroflexi;c__Gitt-GS-136 | NMR | -0.452 | 0.035 |
| | Chloroflexi;c__TK17 | NMR | -0.594 | 0.004 |
| | Verrucomicrobia;c__[Methylacidiphilae] | NMR | 0.506 | 0.016 |
| natural_depth_15-30 | | | | |
| | ;c__ | NMR | -0.579 | 0.030 |
| | Acidobacteria;c__S035 | NMR | -0.560 | 0.037 |
| | Chloroflexi;Other | NMR | -0.632 | 0.015 |
| | Chloroflexi;c__Ellin6529 | NMR | -0.538 | 0.047 |
| | Chloroflexi;c__Gitt-GS-136 | NMR | -0.622 | 0.018 |
| | Gemmatimonadetes;c__Gemm-5 | NMR | -0.554 | 0.040 |
| | Planctomycetes;c__ | NMR | -0.645 | 0.013 |
